# Finding an optimal sequencing strategy to detect short and long genetic variants in a human genome

**DOI:** 10.1101/2025.05.30.656631

**Authors:** Robert J. M. Eveleigh, Sarah J. Reiling, J. Hector Galvez, Mathieu Bourgey, Jiannis Ragoussis, Guillaume Bourque

## Abstract

Advances in DNA sequencing have transformed genomics, enabling comprehensive insights into human genetic variation. While short-read sequencing (SRS) remains dominant due to its high accuracy and affordability, its limitations in complex genomic regions have spurred the adoption of long-read sequencing (LRS) platforms, such as those from Pacific Biosciences (PacBio) and Oxford Nanopore Technologies (ONT). Despite these advances, there is still a lack of systematic, large-scale benchmarking of variant calling performance across diverse platforms, variant types, sequencing depths, and genomic contexts. Here, we present a comprehensive benchmark of sequencing technologies and variant calling algorithms, evaluating their performance in detecting single nucleotide polymorphisms (SNPs), insertions/deletions (indels), and structural variants (SVs). We show that while SRS combined with DeepVariant or DRAGEN offers excellent small variant detection in well-mapped regions, LRS technologies significantly outperform SRS in complex regions and SV detection. PacBio achieves high SNP and smaller SV accuracy even at moderate coverage, while ONT excels in detecting large SVs. The SV callers Dysgu and SVIM emerged as top performers across LRS datasets. Our results highlight that no single platform is optimal for all variant types or regions: SRS remains optimal for high-throughput small variant detection in accessible regions, whereas LRS is critical for capturing SVs and resolving difficult-to-map loci. These findings offer practical guidance for selecting sequencing technologies, coverage and variant calling strategies tailored to specific research or clinical goals, contributing to more accurate and cost-effective genomic analyses.

**Highlights:** ● State-of-the-art short variant calling algorithms are highly comparable but focus on precision and sensitivity differently

● Short-read technologies outperform long-read technologies for small SNP and InDels, with the exception of difficult-to-map variants.

● To capture the majority of variants, a minimum coverage of 15x for PacBio, 20x for SRS, or 30x for ONT is required. However, optimal coverage depends on zygosity, variant type, and the region of interest.

● Long-read technologies outperform short-read technologies for all validation sets tested for deletions and insertions in all size categories.

## Introduction

The evolution of sequencing technologies is generally categorized into three generations, each marked by groundbreaking innovations that have progressively improved the speed, cost-efficiency, and accuracy in the detection of various forms of genomic variation.

First-generation Sanger sequencing^1^, while highly accurate, was laborious and expensive, restricting its application to short DNA fragments and hindering large-scale genomic studies. The advent of second-generation, or Next-Generation Sequencing (NGS)—like Illumina’s sequencing-by-synthesis^1^ (SBS) and, more recently, MGI’s DNA nanoball technology^2^— collectively known as short-read sequencing (SRS), dramatically increased throughput and reduced costs, enabling genome-wide analyses at an unprecedented scale. However, NGS’s reliance on short reads presented challenges, particularly in repetitive genomic regions^3,4^, leading to potential sequencing errors and gaps in genome coverage. To address these limitations, synthetic and third-generation, or linked- and long-read sequencing (LRS) technologies, including 10x Genomics’ linked-read sequencing technology^5^, PacBio’s Single Molecule, Real-Time (SMRT)^6^ sequencing and Oxford Nanopore Technologies’^7^, emerged.

These platforms introduced the ability to reconstruct long-range genomic information or sequence long stretches of DNA in a single read providing comprehensive insights into complex genomic architectures, including repetitive sequences and large structural variants.

Furthermore, LRS offer functionalities like real-time sequencing and epigenetic analysis. While ongoing efforts focus on minimizing error rates and enhancing bioinformatics algorithms to improve data accuracy and utility, third-generation sequencing represents a significant leap forward.

The past decade was marked by advancements in small (SNP and Indels) and large (structural; SV) germline variant detection, driven by improvements in sequencing technology, bioinformatics and computational infrastructure. Early tools like GATK^12^ HaplotypeCaller laid the foundation for detecting single nucleotide polymorphisms (SNPs) and small insertions or deletions (indels) from SRS. More recently, the advent of machine learning (ML)-powered tools such as Clair3^10^, DeepVariant^11^ and Pepper-Margin-DeepVariant^12^ has been coupled with advancements and utilization of hardware acceleration with tools like DRAGEN^14^, Parabricks^15^, MegaBolt^16^ to enable rapid, large-scale variant detection while significantly enhancing sensitivity and accuracy for both SRS and LRS.

Structural variant (SV) —large (typically greater than 50bp) deletions, insertions, duplications, inversions, and translocations— detection has similarly evolved, transitioning from limited resolution with early sequencing methodologies and tools to more accurate detection methods through integration of multiple callers like MetaSV^17^ or modern single-call approaches, such as FPGA-accelerated DRAGEN^14^ and the assembly-based and ML filtering Dysgu^18^. Long-range and long-read technologies have further pushed these boundaries, enabling accurate identification of larger and more complex SVs with novel complementary tools like LongRanger^5^, CuteSV^19^, Nanocaller^20^, pbsv^21^, Sniffles2^22^, and SVIM^23^.

As sequencing technologies and variant calling algorithms evolve, robust validation and benchmarking methodologies have become essential for assessing the accuracy, precision, and reliability of genomic data^24,25^. Organizations like the Genome in a Bottle^26^ (GIAB) consortium, SEQC2^27^ and the Global Alliance for Genomics and Health^24^ (GA4GH) have played pivotal roles in developing standards for benchmarking, and providing high-confidence reference datasets that allow for the detection and quantification of errors, biases, and inconsistencies in genomic analyses. GIAB has established itself as a leader in creating well-characterized reference genomes with comprehensive, high-confidence variant calls, including small variants^28^ (SNVs and indels), structural variants^29,30^ (SV), tandem repeats^31^, and preliminary large and small variant derived from near telomere-to-telomere (T2T) assemblies. The GIAB reference genome panel includes widely used samples such as HG001 (Centre d’Etude du Polymorphisme Humain (CEPH) NA12878), the Ashkenazim trio (HG002, HG003, and HG004), and the Han Chinese Trio (HG005, HG006, and HG007)^32^. This population-diverse genome panel promotes equity while providing key benchmark standards for high-confidence variant calling, and can be instrumental in validating Mendelian inheritance patterns and *de novo* variant calls.

The recently released GIAB small variant dataset (version 4.2.1) and structural variant (SV) dataset (version 0.6) build upon earlier versions by integrating data from diverse populations, incorporating new sequencing technologies, and leveraging multiple variant calling algorithms. This comprehensive approach mitigates platform-specific limitations and enhances variant detection by expanding coverage across the reference genome—reaching 92% coverage of the GRCh38 autosomes, up from 85% in the previous release (v3.3.2). As a result, approximately 300,000 additional SNVs and 50,000 indels were identified, largely due to the inclusion of challenging genomic regions such as segmental duplications, self-chains, and the MHC region—areas historically difficult to resolve with SRS. This also included 16% more exon variants related to challenging, clinically relevant genes providing a more robust resource focusing on curating challenging medically relevant genomic regions^33^ (CMRG), which include loci frequently implicated in genetic disorders, pharmacogenomics, and cancer. This clinical emphasis ensures that the dataset is particularly valuable for validating diagnostic assays targeting these areas. Furthermore, the dataset includes an expanded and curated collection of structural variants to provide a robust benchmark for detecting large insertions, deletions, and other complex variants e.g. multiple variants within 10bp of each other.

Complementing GIAB’s efforts, GA4GH^34^, have contributed by stratifying genomic regions to enhance benchmarking assessments. These stratified regions delineate areas of the genome with specific characteristics, such as coding regions, low mappability regions, high GC content regions, various types of repetitive regions, or medical relevance, enabling researchers to evaluate sequencing technologies and algorithm performance under diverse genomic contexts. To support this, small variant benchmarking tools such as vcfeval^35^, hap.py^24^, and large variant (SV) tools like truvari^36^ are widely employed. These tools not only provide robust methodologies for comparing variant calls against high-confidence reference datasets to generation of metrics like precision, recall, and F1-score but also enable stratification of variants to facilitate more nuances assessments. Their integration into benchmarking workflows ensures a rigorous and standardized evaluation of variant calling performance, allowing researchers to gain insights into the accuracy and limitations of sequencing platforms and bioinformatics pipelines.

In light of these advances, there has been an increase in analyses identifying the strengths and weaknesses of sequencing technologies^28^ and bioinformatic tools^20^. However, there is limited information comprehensively summarizing the benefits and limitations of different sequencing technologies and variant callers across the full spectrum of variants (SNVs, indels and SVs), for different sequencing coverage and genomic contexts. In this article, we evaluate recent sequencing technologies using the latest validation truth sets from the Ashkenazim trio, specifically HG002, to delineate the strengths and weaknesses of state-of-the-art small and large variant callers using the latest benchmarking methodologies. Our analysis evaluates data under different scenarios, including variant type, genomic region, zygosity, and sequencing coverage. With this information, we provide recommendations for the optimal combination of sequencing technology, coverage, and variant caller to best suit specific research and clinical scenarios.

## Results

### Short-read variant calling algorithms prioritize precision and sensitivity differently

To evaluate the accuracy of short and long read sequencing techniques, we assessed various libraries using the HG002 cell line (**<u>Methods</u>**). Specifically, we generated libraries and sequenced this cell line using four different platforms: two SRS, (Illumina Xten and MGI G400), one synthetic 10X Genomics (10X), and one LRS technology, Oxford Nanopore Technologies (ONT). Additionally, we obtained one Pacific Biosciences HiFi (PacBio) LRS dataset (PB) from GIAB directly. The average depth of coverage obtained for each platform, as well as the average physical molecule length are summarized in **<u>Table S1</u>**. These libraries allowed us to test the performance of read-length on the detection of single nucleotide polymorphisms (SNPs), small (<50bp) indels and large (>50 bp) insertions and deletions using the latest GIAB (4.2.1 and 0.6 SV) and challenging, medically-relevant genes (CMRG) validation sets to generate precision, recall and F1-score metrics.

First, we tested the following variant detection tools on the two SRS technologies: four CPU-based callers: GATK3, GATK4 haplotype caller, DeepVariant, Clair3, two graphics processing units (GPU)-based: Parabricks and MegaBOLT, and one field-programmable gate array technology (FPGA)-based: DRAGEN using default settings (see **<u>Methods</u>**). To evaluate small variant performance, we identified all easy- and difficult-to-map variants within each validation set to generate benchmarking metrics. All algorithms achieved an F1-score above 99.3% and 95.3% for SNPs, and above 98.2% and 91.0% for indels within the GIAB and CMRG validation sets, respectively (**<u>Table S2</u>**). DeepVariant ranked the highest across both variant types and validation sets, based on F1-score, followed closely by DRAGEN, then by Clair3, GATK4, GATK3, Parabrick, and Megabolt.

Although F1 scores varied slightly, MGI exhibited lower false discovery (FDR) and false negative rates (FNR) for the GIAB dataset, while Illumina (ILL) showed the inverse in the CMRG dataset. Despite this difference, the relative ranking of variant calling algorithms remained consistent between both SRS technologies (**Figure 1a**). Therefore we focused on ILL data to compare the subtle differences between the tools by assessing the performance of each variant calling algorithm. By comparing FDR and FNR, we observed small differences in the tools’ abilities to control type I and type II errors. DRAGEN was the most sensitive of all algorithms tested, with an FNR measure of 0.73% and 5.38% for GIAB and CMRG, respectively with DeepVariant following closely with a slight increase in FNR to 0.77% and 5.59%. Conversely, DeepVariant had the lowest FDRs values of 0.015% and 1.22%, halving the number of false positives when compared to DRAGEN (0.038% and 2.68%). The other ML-based caller Clair3 had lower FDR (0.51% and 3.33%) than all GATK-based caller GATK4 (0.61% and 3.35%), MegaBOLT (0.64% and 3.12%), Parabricks (0.64% and 3.34%) but had an FNR (0.96% and 6.68%) between the lower FNR CPU-based GATK caller (GATK4: 0.88%, 6.14% and GATK3: 0.85%, 5.96%) and GPU-based callers (MegaBOLT: 1.08%, 6.55% and ParaBrick: 0.95%, 6.65%). Among the GATK-based callers, the CPU-based callers outperformed the GPU-based callers with GATK4 attaining a lower FDR than GATK3 (0.61% vs 0.7%) but GATK3 had lower FNR than GATK4 (0.85% vs 0.88%). MegaBOLT and Parabricks, each utilizing the GATK4 framework, both had similar performances to GATK4. However, Megabolt (1.08% and 6.55%) had slightly larger FNR than Parabricks (0.95% and 6.3%), while Parabricks (3.34% vs 3.12%) had slightly larger FDR for CMGR when compared against Megabolt.

**Figure 1.**
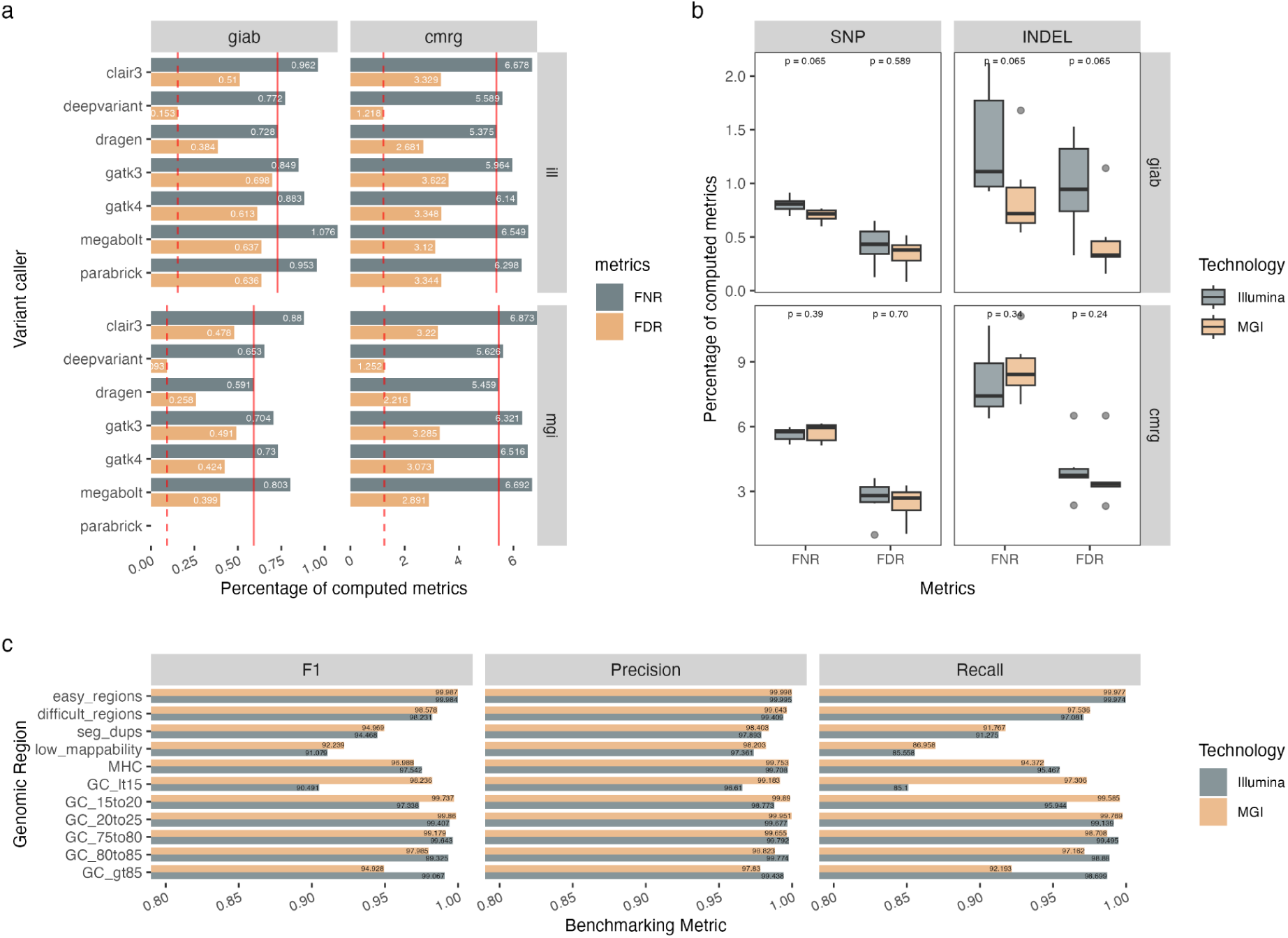
- SRS Variant Caller Priorities: Precision vs Sensitivity a. Comparing the percentage false negative (FNR) and false discovery (FDR) rates for both GIAB (4.2.1) and CMGR validation sets across diploid germline callers using one lane of 41x Illumina Xten and 45x MGI G400 sequencing data. Each label represents either the FN or FD rates with a dash or solid vertical line to represent minimal values representing the top ranked performer. b. Comparing the FNR and FDR distributions for Illumina and MGI technologies across multiple germline variant callers for SNPs, indels and both validation datasets. Two-sided Wilcox tests were performed to identify that the distribution of FNR and FDR are not significantly different (<p-value 0.05) for either validate set or variant type. c. Barplot depicting the F1 score, recall and precision differences for Illumina and MGI technologies across various genomic contexts. MGI has higher F1 scores for difficult-to-map regions which include MHC, segmental duplications and low mappability regions as well as regions with high GC context (>70%). Conversely, ILL has higher F1 score in AT-rich regions (>70%).

Although the ranking of the variant callers were similar between both SRS technologies (**Figure 1a**), we further stratified the SRS data by variant type (SNPs and indels), validation set (GIAB and CMGR) and genomic context according to GIAB stratified regions to identify inter-technology differences. First, we analyzed the FNR and FDR distributions across all callers (excluding Parabricks) by variant type and validation set using a two-sided Wilcoxon test (significance threshold: p=0.05). For SNPs, the distributions were not significantly different across the two validation sets. In contrast, for indels, differences were observed in the GIAB dataset, suggesting that MGI may outperform Illumina in detecting these variant types. This improved performance may be attributed to MGI’s larger insert size (378 bp vs. 308 bp; **Figure 1b**). Next, we stratified the variant calls made by DeepVariant according to GIAB annotated genomic regions to assess inter-technology differences in specific genomic contexts. MGI demonstrated marginally better F1 scores in difficult-to-map regions, driven by increased sensitivity in segmental duplications, low-mappability regions, and AT-rich regions (>75%). Conversely, Illumina outperformed MGI in the MHC region and GC-rich regions (>75%) (**Figure 1c**).

In addition to overall performance in small variant detection, we also compared the computational efficiency, including walltime (i.e. total running time), for each of the six short-read variant callers. As anticipated, callers utilizing hardware acceleration, such as FPGA or GPU, as is the case with DRAGEN, MegaBOLT, and Parabricks, were orders of magnitude faster than CPU-based callers (**<u>Table S3</u>**). Our assessment of total walltimes across SRS technologies using CPU-based callers showed that MGI data generally took longer to process than ILL (approximately - ILL: 33.5hr±2hr and MGI:46hr±4hr), which is likely due to higher coverage.

Additionally, the GATK4 haplotype caller took almost twice as long (7h05min versus 4h32min) to run than GATK3 for MGI data, but the inverse was observed in the Illumina dataset. This suggests that GATK4 code could be internally optimized for ILL data.

### SRS technologies outperform LRS technologies for small variant detection but LRS is better in certain genomic regions

To assess the overall performance between SRS and LRS technologies, we compared the different sequencing technologies and variant calling algorithms using standardized practices developed by the GA4GH benchmarking team and the GIAB consortium. Evaluating the top performers for all small events (including both SNPs and indels) and across all selected algorithms and technologies, we observed that all technologies, with the exception of ONT, all achieved an overall F1-score greater than 97% (**<u>Table S4</u>**). However, by masking variants in proximity to homopolymers; known regions of sequencing bias and error^20^, the Clair3 algorithm using ONT data achieved an overall F1-score greater than 97.0% (**<u>Figure S2</u>**). DeepVariant, applied to SRS data (MGI: 99.6% and ILL: 99.5%), outranked all other callers and technologies, followed by Clair3 with PacBio (98.7%), LongRanger with 10X (97.8%) and Clair3 with ONT (97.0%). The remaining ONT callers: Nanocaller and Pepper were the least performing. GATK4 had varying performance depending on the sequencing technology: SRS performed better (MGI: 99.42% and ILL: 99.25%) than PacBio (87.27%).

When segregating the results by variant type (**Figure 2**), all technologies achieved an F1-score greater than 99% for SNPs, with PacBio (using Clair3: 99.92%) outperforming DeepVariant in SRS (MGI: 99.62% and ILL: 99.56%), followed by 10X (99.21%) and finally, ONT (99.16%). However, when comparing indels, LRS and synthetic technologies had lower F1-scores (PB 91.34%, ONT 72.49% and 10X 88.51%). Conversely, SRS technologies maintain similar performance with indels (MGI: 99.63% and ILL: 99.37%) than with SNPs. In sum, overall SRS outperform LRS and synthetic technologies in terms of F1-scores when both variant types are combined, but separately LRS outperform SRS and synthetic technologies for SNP and vice versa for indels.

**Figure 2.**
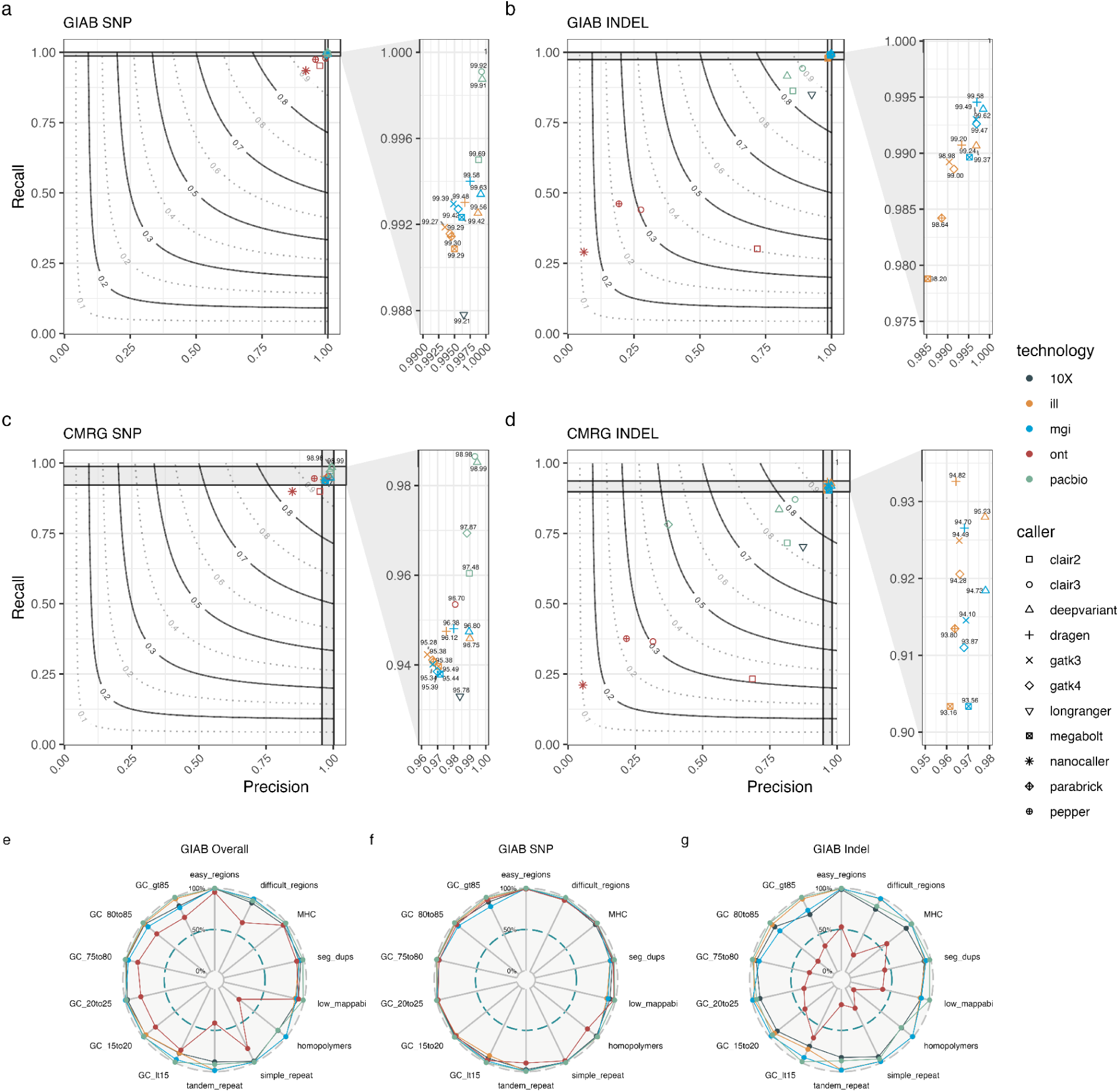
- Comparative Performance of SRS and LRS for Small Variant Detection Precision-Recall graph with F1-score contours were generated for SNPs and Indels using two validation sets: GIAB 4.2.1 (a,b) and CMRG (c,d) for five technologies: 10X (black), Illumina (yellow), MGI (blue), ONT (red) and PacBio (green). F1 scores across all tested technologies: SRS (Illumina and MGI), LRS (PacBio and ONT) and synthetic (10X) stratified by GA4GH easy and difficult to map regions for e) GIAB dataset for both SNPs and Indels combined, f) GIAB SNPs only g) GIAB Indels only

To assess the impact of different genomic regions on variant calling performance, we compared Clair3, one of the top-performing variant callers across most technologies, with 10X LongRanger by stratifying each validation set across GIAB’s easy-to-map and difficult-to-map regions (**Figure 2**, **<u>Figure S3</u>**). As expected, the F1 scores for easy-to-map regions were comparable across all technologies and validation datasets, with the exception of indels using ONT, which continued to underperform. LRS technologies generally outperformed SRS, particularly for SNPs (both PacBio and ONT) and indels (PacBio only) in regions associated with mapping challenges, such as segmental duplications and low-mappability areas. However, ONT exhibited reduced performance near repeats (simple and tandem) and homopolymer regions, likely due to known sequencing biases. PacBio demonstrated more consistent performance across high (≥75%) GC- and AT-rich regions, while MGI and Illumina showed reductions in F1 scores in GC-rich and AT-rich regions, respectively. Interestingly, although the synthetic 10X technology slightly underperformed in easy-to-map regions compared with SRS, it showed marginally better performance in difficult-to-map regions, highlighting its potential for resolving complex genomic features.

### A minimum of 20X average whole-genome depth of coverage is required to capture the majority of variants

To determine the minimum depth of coverage needed to capture most small SNPs and indels, samples from the Ashkenazim son HG002 (GM24385), paternal (HG003) and maternal (HG004) individuals (**<u>Figure S4</u>**) were downsampled to various mean depths (ranging from 2x to 120x depending on the technology; see **<u>Methods</u>**). These samples, from both SRS (ILL and MGI) and LRS (ONT2 and PB2) sequencing, were then analyzed using the Clair3 variant caller.

Performance was benchmarked against both GIAB and CMRG truth sets, with results categorized into GA4GH easy-to-map and difficult-to-map regions. Analysis of the validate truth sets for GIAB and CMRG revealed that the majority of true positive SNPs (over 70%) were categorized as residing in easy-to-map regions, whereas the majority of true positive indels (over 67%) were categorized as residing in difficult-to-map regions (**<u>Table S2</u>**).

In easy-to-map regions, SNP and indels variant discovery increases with increasing depth: in other words, there is an increase in true positive events and a decrease in the number of false positive events as depth increases, with diminishing returns after 15-fold for PB, 20-fold for ILL and MGI and 30-fold coverage for ONT (**Figure 3a**). In difficult-to-map regions, a similar trend was observed. However, most sequencing technologies require deeper sequencing to reach their respective plateau with ILL requiring at least 60x coverage, ONT at least 45x and PacBio requiring similar sequencing depth (25-fold) as observed with the easy-to-map regions.

**Figure 3.**
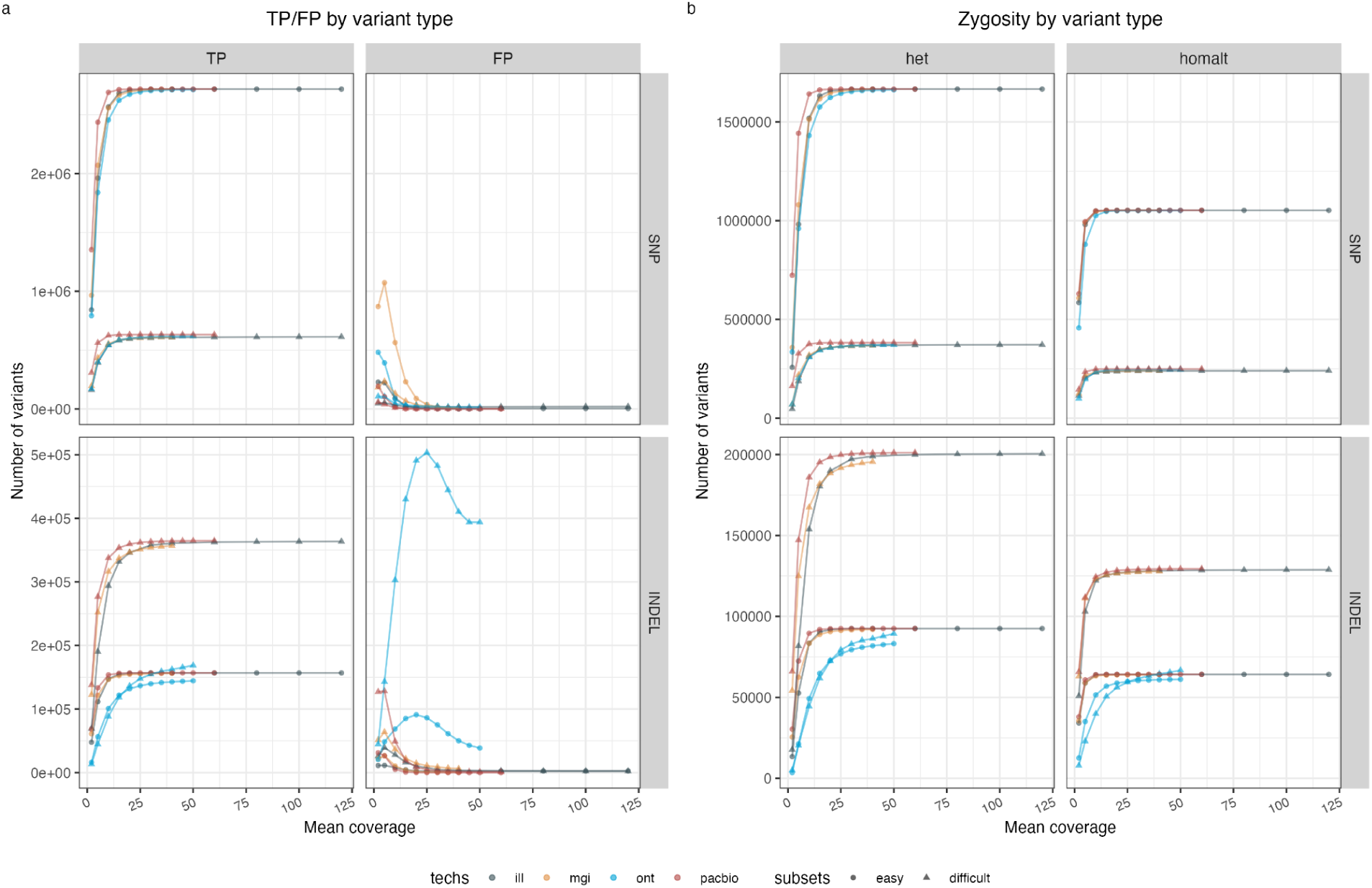
- Optimal Small Variant Detection Requires at least 20 fold Whole Genome Coverage a. Coverage titration of HG002 sample using four technologies: Illumina (ILL) downsampled from 2x to 120x mean coverage, MGI 2x to 45x, ONT 2x to 50x, and PacBio 2x to 60x. True positive (TP), and false positive (FP) counts were collected for both easy-to-map and difficult-to-map regions for SNPs and InDels within the full GIAB and CMRG validation set. Total percentage of true variants were labelled on at each coverage point. b. Line plots showing the proportion of true positive SNP and InDel events detected at different mean depths of coverage for ILL data from the GIAB HG002 in both easy- and difficult regions to-map regions of both validation sets (GIAB 4.2.1 and CMRG). Results are stratified by homozygous events (yellow) and heterozygous events (grey).

When investigating the impact of varying input depth on zygosity detection, heterozygous (het) calls were identified more frequently in both truth sets, constituting more than 60% of true variants. Among the easy-to-map regions, identifying 99% or more of homozygous alternative (homalt) SNPs and indel positions requiring less coverage than for het variants. Illumina and PacBio technologies both required 10x coverage to call over 99.6% of the true positive homalt SNPs and indels, and 20x and 15x respectively to capture more than 99.3% of the het calls.

ONT required 15x (99.5%) and 30x (99.2%) coverage to capture the majority of true homozygous and heterozygous SNPs, but could not detect more than 88.6% het and 95.5% homalt InDel calls at 50x coverage (**Figure 3b**). In the difficult-to-map regions, PacBio was the only technology able to capture more than 99% of both zygosity types, requiring 15x (99.3% for hets) and 10x (99.4% for homalts) coverage for SNPs and 25x for het and homalt indels.

Illumina at maximum coverage of 120x detected only 96.9% and 96.4% of difficult-to-map SNPs, but was able to identify over 99% of indels at 60x and above. Lastly, ONT identified a maximum of 96.9% and 98.5% of heterozygous and homozygous SNPs and less 51.4% of the known indels were identified at 50x.

### Long-Read technologies outperform Short-Read technologies in Structural Variant detection

Next, we evaluated the performance of the various sequencing technologies in detecting structural variants (SVs). The performance of different SV calling tools was assessed by using svbench to generate standardized metrics across three validation datasets: (1) the full Tier 1+2 dataset, (2) a high-confidence Tier-1 subset, and (3) a subset focused on clinically relevant genes (CMRG). The Tier 1+2 dataset includes 15,001 deletions (DEL) and 15,948 insertions (INS) over 30 bp (excluding 4,413 DEL and 4,141 INS below 30 bp) from GIAB’s HG002 sample. It combines well-supported variants from multiple technologies (Tier-1) with less confident variants useful for exploring detection challenges (Tier-2). The Tier-1 subset contains a quarter of known HG002 SVs, with 4,117 DEL and 5,261 INS over 50 bp, all sequence-resolved. The CMRG subset focuses on 89 high-confidence, sequence-resolved DEL and 105 INS near 273 clinically relevant genes (**<u>Table S6</u>**).

Comparative analysis of SV calling methods across sequencing technologies was performed using two metrics: discovery F1 reflecting the balance of precision and recall for correctly identifying the presence and relative location of structural variants, regardless of genotype and genotyping (GT) F1 to measure the accuracy of both detecting and correctly genotyping the variant; this metrics provides a more stringent assessment of SV caller performance.

Assessment using these metrics revealed that LRS consistently outperformed SRS and synthetic technologies across all validation sets and variant types, as evidenced by substantially higher discovery F1 and GT F1 (**Figure 4**, **<u>Figure S5</u>**). Specifically, LRS achieved average discovery F1 of 73.0%, 96.3%, and 94.4% for DEL in the Tier1+2, Tier1, and CMRG datasets, respectively —1.23 to 1.68 times higher than SRS and 1.35 to 1.84 times higher than synthetic technologies. The difference was even more pronounced for INS, where LRS attained average discovery F1 of 74.4%, 94.6%, and 86.6%, outperforming SRS by 2.09 to 2.63 times, while 10X failed to detect any INS variants (**<u>Table S6</u>**). LRS also demonstrated significantly improved genotyping accuracy. For DEL, LRS achieved GT F1 scores of 49.5%, 94.6%, and 77.9% for Tier1+2, Tier1, and CMRG, respectively, reinforcing that genotyping variants is generally more challenging, particularly in the more complex Tier1+2 and CMRG datasets. GT F1 was up to 1.82 times lower in SRS and 2.79 times lower in 10X compared to LRS. Similarly, for INS, LRS GT F1 scores were 51.0%, 90.3%, and 68.3%, exceeding the SRS average by up to 3.17 times.

**Figure 4.**
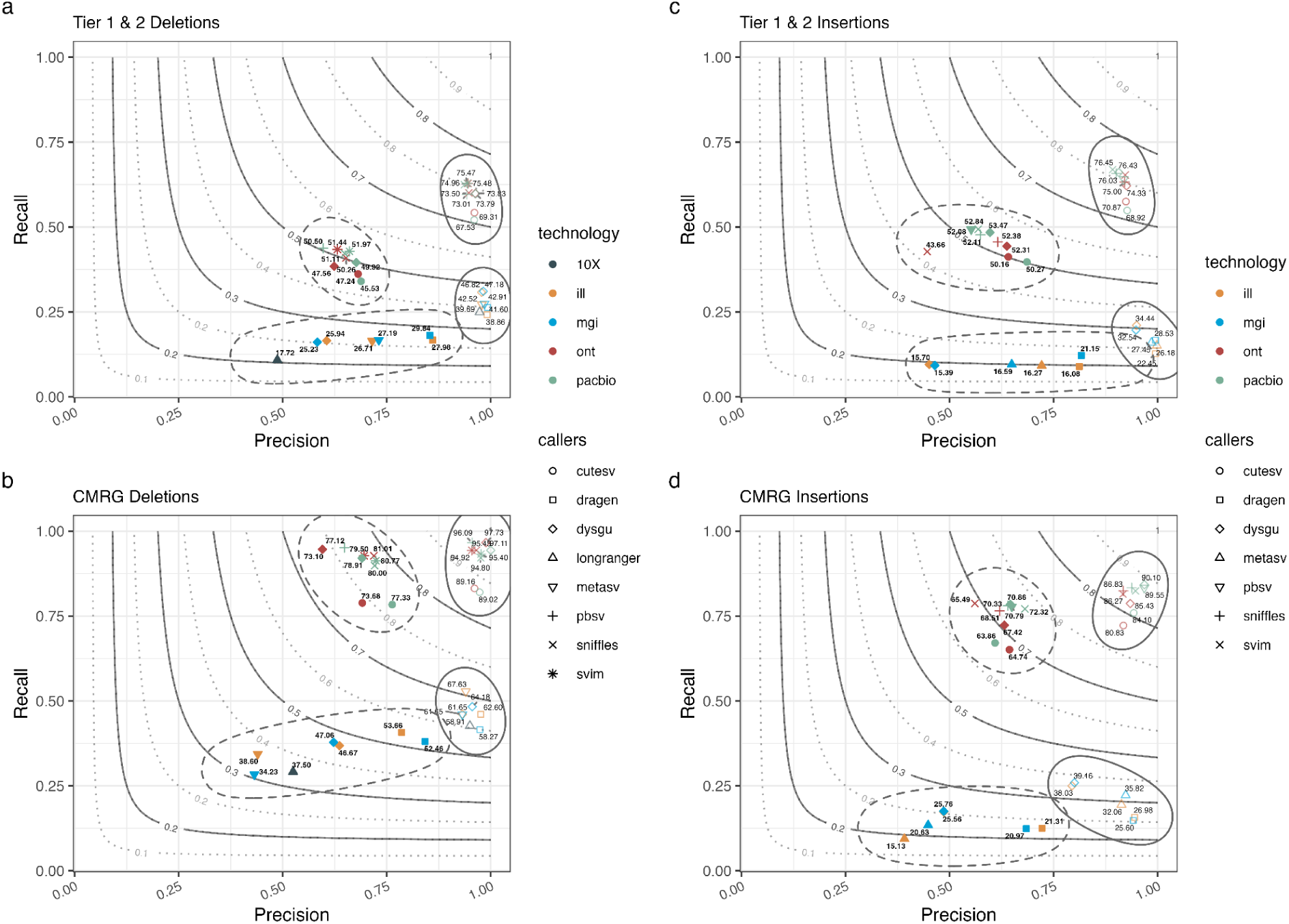
- Comparative SV Detection: LRS vs. SRS and Synthetic Precision-recall curves across the two complex validation sets: Tier 1+2 contains approximately 15k deletions a) and insertions c) over 30bp, Tier 1 contain 4117 and 5261 high quality deletions b) and insertions d), and CMRG containing 89 and 105 deletions c) and insertions f) in 273 medically relevant genes. These three datasets were used to interrogate five different sequencing technologies and eight different SV callers.

### Structural variant calling performance vary depending on technology, variant type and validation set

Within LRS technologies, discovery and GT F1 performance varied by technology, SV type and validation set. ONT performed slightly better in DEL discovery, while PB typically excelled at identifying INS (**Figure 4**, **<u>Figure S5</u>**). However, this pattern was not observed for GT F1 with PB generally outperformed ONT across both variant types and most validations sets. More specifically, for DEL, in the more complex Tier 1+2 dataset, SVIM achieved the highest discovery F1 for both technologies (ONT: 75.48%; PB: 75.47%), with PB SVIM identifying the highest number of true positives (recall: 62.88%), narrowly surpassing ONT SVIM (62.81%).

However, ONT Dysgu led across both high-confidence datasets (Tier 1: 97.72%, CMRG: 97.69%), slightly outperforming PB pbsv (97.57% and 96.09%, respectively). In contrast, GT F1 largely favored PB, with PB SVIM achieving the highest GT F1 for DEL in Tier 1+2 (51.97%) and Tier 1 (96.78%), while ONT Sniffles2 (82.35%) led in the CMRG dataset. For INS, again SVIM had the highest discovery F1 for both technologies in Tier 1+2 (PB: 76.45%, ONT: 76.45%), with PB achieving the highest recall and ONT the highest precision. PB Dysgu consistently outperformed other PB callers in discovery (Tier 1: 96.52%, CMRG: 90.1%), exceeding the top ONT performers—SVIM (Tier 1: 95.91%) and Sniffles2 (CMRG: 86.83%). Similarly, for GT F1, PB maintained dominance, with Dysgu leading in Tier 1+2 (53.47%) and Tier 1 (93.23%), outperforming ONT Sniffles2 (52.38% and 92.62%). In the CMRG dataset, PB SVIM (72.32%) surpassed ONT Sniffles2 (70.33%).

Similar to LRS, the performance of SRS and synthetic technologies varied: MGI generally showed better DEL discovery, while ILL had a slight edge in INS particularly for Tier 1+2 and Tier 1, a trend that reversed in the CMRG dataset. SRS GT F1 followed this trend, with MGI outperforming ILL for DEL and vice versa for INS unlike in LRS. More specifically, in terms of discovery F1 for DEL, Dysgu led in Tier 1+2 (MGI: 47.18%; ILL: 46.82%), and MGI DRAGEN in Tier 1 (80.17%), surpassing ILL MetaSV (79.25%). However, ILL MetaSV outperformed MGI Dysgu in CMRG (67.63% vs. 64.18%). For GT F1 of DEL, MGI outperformed ILL across most datasets, except in CMRG where ILL DRAGEN excelled. Specifically, DRAGEN achieved the highest GT F1 in Tier 1+2 (MGI: 29.84%; ILL: 27.98%) and Tier 1 (MGI: 79.11%; ILL: 75.73%), but ILL DRAGEN led in CMRG. Moving to INS, ILL Dysgu showed the highest discovery F1 in Tier 1+2 (ILL: 34.44%, MGI: 32.54%), while in Tier 1, ILL Dysgu (52.74%) and MGI MetaSV (51.23%) were top performers. In CMRG, MGI Dysgu (39.16%) slightly exceeded ILL Dysgu (38.03%). For GT F1 of INS, MGI DRAGEN (21.15%), MetaSV (41.95%), and Dysgu (25.67%) were the top performers in Tier 1+2, Tier 1, and CMRG, respectively. ILL MetaSV excelled in Tier 1+2 (16.27%) and Tier 1 (40.1%), and ILL Dysgu led in CMRG (25.56%). Notably, 10X LongRanger consistently underperformed in both discovery and GT F1 for DEL (discovery: Tier 1+2: 39.69%, Tier 1: 71.15%, CMRG: 58.27%; GT F1: Tier 1+2: 17.72%, Tier 1: 49.62%, CMRG: 37.5%), and failed to detect any insertions, reinforcing its limited efficacy compared to ILL and MGI.

### Size-stratified performance of SV callers varies across sequencing technologies

To gain a more granular understanding of each technology’s performance given various SV callers, we stratified the caller results into bins of varying, non-overlapping sizes, considering known peaks associated with Alu elements (±300 bp) and full-length LINE1 elements (±6000 bp; Zook et al., 2020). Specifically, we created bins for SV sizes ranging from 30-50 bp (exclusively in the Tier 1+2 dataset), 51-250 bp, 251-500 bp, 501-6000 bp, and greater than 6000 bp.

Stratification of the three validation datasets revealed a general trend: the abundance of both deletions (DEL) and insertions (INS) decreased as SV size increased. Notably, the 51-250 bp size range exhibited the highest number of SV events, with a subsequent decline in event counts observed in larger size bins (**Figure 5**). This pattern was consistently observed in the high-confidence datasets, Tier 1 and CMRG, with CMRG lacking events greater than 6000bp. However, the comprehensive whole-genome dataset, Tier 1+2, presented a slight deviation, with the 30-50 bp size range showing the second highest abundance of SV events (**<u>Table S6</u>**).

**Figure 5.**
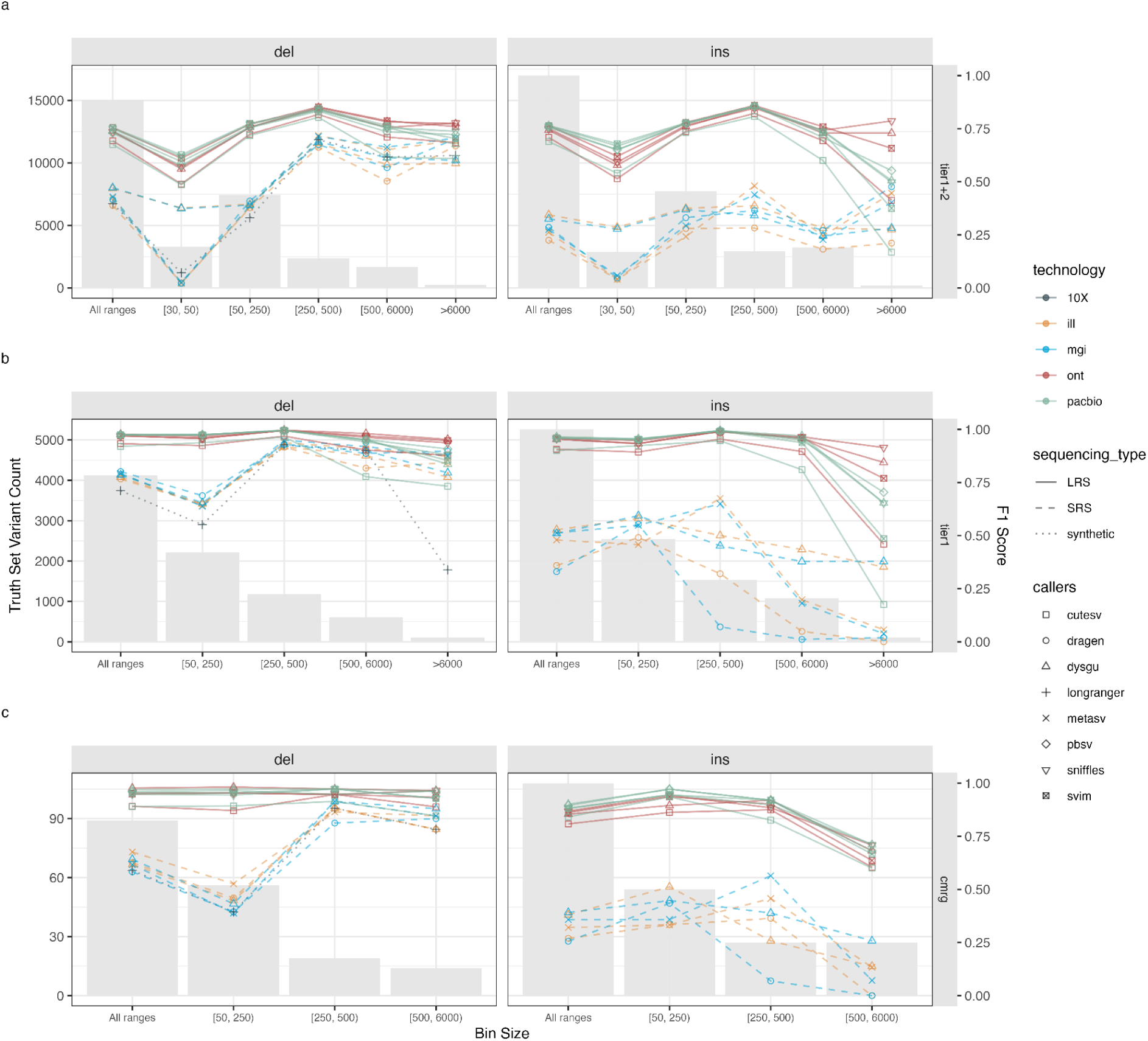
- Technology-Dependent SV Caller Performance Across Sizes Performance of each technology: LRS, SRS and synthetic described as the median percent of all discovered true positive events across all structural callers tested and stratified by SV sizes of 30 to 50 bp (Tier 1+2 only), 51 to 250bp, 251 to 500bp, 501 to 6000bp and greater than 6kb. Deletion events for truth set a) Tier 1+2, b) Tier 1 and c) CMRG and insertion events for d) Tier 1+2, e) Tier 1 and f) CMRG

Applying our binning strategy across all validation sets, SV types, and technologies reinforced a key observation: LRS technologies consistently detected a greater number of both DEL and INS compared to SRS and synthetic data for all validation sets and sv sizes. This analysis highlighted a higher abundance of LRS-detected SV events within the smaller size ranges for both DEL and INS—specifically in the 30–50 bp (Tier 1+2 only) and 51–250 bp bins—as well as for larger INS events exceeding 500 bp. In these bins, LRS technologies identified approximately 3.81 times more DEL events in the smallest Tier 1+2 range (30–50 bp), and 1.9 to 3.12 times more DELs in the 51–250 bp range, depending on the validation set (**<u>Table S7</u>**).

Similarly, LRS identified 6.64 times more INS events in the Tier 1+2 30–50 bp bin, and between 2.48 and 3.51 times more INS in the 51–250 bp range. At the opposite end of the size spectrum, LRS technologies detected between 4.2 and 9.78 times as many INS events in the 500–6000 bp range, and 1.77 times (Tier 1+2) to 5.94 times (Tier 1) more events larger than 6000 bp. While the synthetic 10X data underperformed relative to LRS, it detected slightly more events than SRS in specific size ranges—the 250–500 bp (Tier 1+2) and 500–6000 bp ranges (both Tier 1+2 and Tier 1).

When comparing the two LRS technologies, we observed that PB typically achieved slightly higher F1 accuracy for SVs smaller than 250 bp, while ONT achieved higher accuracy for regions larger than 250 bp which was most pronounced for larger insertions (>6000bp).

However, in some instances, performance varied based on the validation set, SV type and SV size, most notably in Tier 1 INS where PB identified more 250-500bp events. Among LRS SV callers, SVIM generally performed best across most regions in both Tier 1+2 DEL and INS with the exception of events larger (>6 kb), where Sniffles2 and pbsv had the highest F1 score for ONT and PB, respectively. For the high-confidence datasets, Dysgu was the most frequent LRS top performer in Tier 1 DEL with pbsv (50-250 and >6 kb) and SVIM (250-500bp) being the top performers in specific size ranges for PB. For Tier 1 INS, SVIM (50-250bp) and dysgu (251-500bp) were the ONT top-performers in for SV <500bp with Sniffiles2 outperformed other callers for SV larger than 500bp. Dysgu was the most frequent top-performer for PB Tier 1 INS with pbsv performing best in 250-500 and >6 kb regions. For CMRG, Dysgu had the highest F1 score across all ranges for DEL and PB INS, but for ONT INS Sniffles2 was the top-performer in 50-250 and 500-6000bp ranges. It is important to note that due to low event counts in the CMRG dataset—particularly for larger SVs—Dysgu, SVIM, Sniffles2, and pbsv (PB) often tied as the top performers in these categories.

When comparing SRS technologies and synthetic data, MGI generally outperformed Illumina (ILL) and synthetic data across various size ranges, although exceptions occurred based on validation sets, SV type, and size. Specifically, ILL achieved the highest F1 scores for 250-500 bp INS in Tier 1 and Tier 1+2 datasets, as well as for 500-6000 bp INS in Tier 1, and for 30-50 bp DEL in Tier 1+2, and two size ranges (50-250 bp and 250-500 bp) in the CMRG dataset.

Synthetic data slightly surpassed MGI for 250-500 bp and 500-6000 bp DEL in Tier 1+2. Among SRS and synthetic SV callers, Dysgu was the most frequent top performer, demonstrating the highest INS F1 scores across most ranges and validation sets. However, MetaSV consistently performed best in the 250-500 bp INS range across all datasets. For DEL, the top performer varied. MetaSV typically excelled for Tier 1+2 and Tier 1 DEL events larger than 250 bp, while Dysgu or Dragen (in MGI) performed best for smaller events. In the CMRG dataset, Dysgu was the top performer across all SV sizes for MGI, whereas MetaSV was preferred for ILL, with Dysgu excelling in the 250-500 bp range.

### Minimum coverage requirements for SV discovery vary by technology, validation set and SV type

To determine the minimum sequencing coverage necessary for comprehensive SV detection, we analyzed downsampled datasets. For ILL, we used Dysgu on previously downsampled data, ranging from 2x to 120x coverage, across three validation sets. For long-read sequencing (LRS), we examined two deeply sequenced GIAB datasets: ultralong ONT (ONT2), downsampled from 2x to 50x, and PB Sequel II (PB2), downsampled from 2x to 60x. Consistent with small variant detection, SV discovery improves with increased sequencing depth (**Figure 6**). Specifically, the number of true positive events increases, while false positive events decrease, as depth increases. However, this improvement exhibits diminishing returns beyond 35x coverage for ONT, 40x for PB, and 80x for ILL. This trend holds across different validation sets and SV types. Nevertheless, the optimal SV caller and required coverage vary depending on the validation set and SV type. For SRS, achieving the highest discovery F1 score generally requires a minimum of 80x coverage across most datasets and SV types, with the exception of DEL in the CMRG dataset. However, maximizing true positive (TP) events necessitates specific coverage adjustments. For instance, 120x coverage is required for Tier 1 INS to achieve the highest recall, and 60x coverage is needed for CMRG DEL to maximize both F1 and recall.

**Figure 6.**
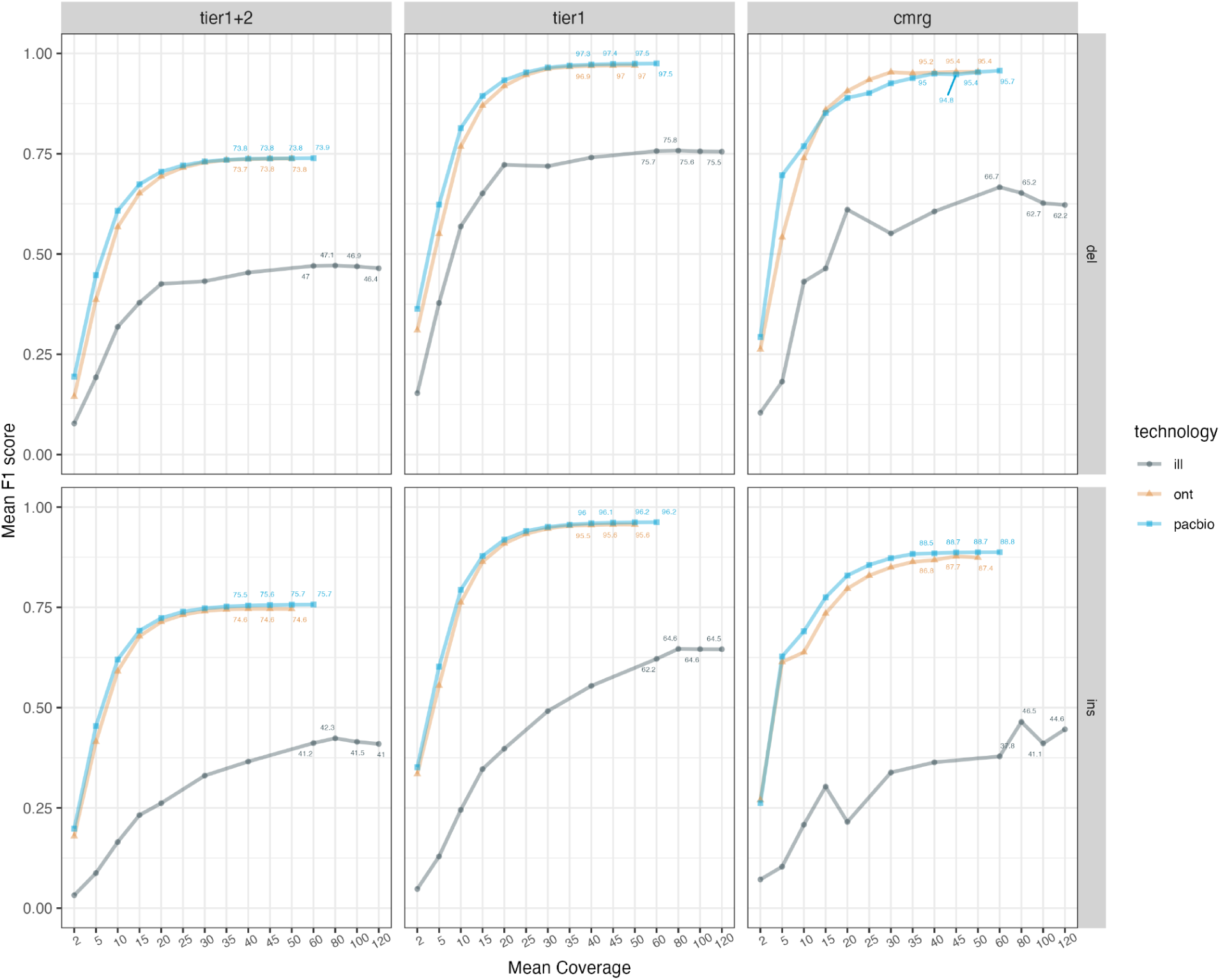
- SV discovery: Coverage Varies by Technology, Validation, and Type Coverage titration of one SRS technology - Illumina (ill) from 2x to 120x for three lanes of HG002 sequenced on HiSeq X, and two LRS technologies ONT 2 to 50x and PacBio 2x to 60x. Mean F1 scores were collected for both easy to resolve SVs (Tier 1) and challenging SV (Tier 1+2 and CMRG) for DEL and INS within the full GIAB validation set. Labels were added for LRS if means coverage was above 40x and for ILL if it was above 50x

Similarly, for LRS, the highest attainable coverage in this experiment typically yields the highest discovery F1 score due to improved precision with increasing coverage. However, lower coverage suffices for certain SV callers, validation sets, and SV types. Notably, for ONT-called INS using SVIM on the Tier 1+2 dataset, 25x coverage achieves the highest F1 score, and 20x the highest recall. Dysgu required 15x PB data to attain the highest F1 and recall for INS in the CMRG dataset.

## Discussion

In this study, we assessed the performance of various sequencing technologies and variant calling algorithms across a range of variant types, including SNPs, indels, and SVs. Moreover, by utilizing complex and clinical validation datasets across these variant types and by further subsetting using different genomic contexts, we highlighted distinct performance patterns between LRS, SRS, and synthetic technologies, as well as between different variant calling algorithms and their performance at varying coverage depths. These findings provide a nuanced understanding of how technology and methodology choices influence variant detection accuracy, offering insights for optimizing variant calling under certain scenarios (**Table 1**).

**Table 1.** - Top Technologies for Variant Detection by Genomic Region and Validation Set Performance assessment of various technologies for small and large variant detection by determining the top three rankings within specified genomic regions and validation sets

| Region/Validation set | Subregion | Top 1 | Top 2 | Top 3 |
| --- | --- | --- | --- | --- |
| Easy-to-map | SNP | PacBio | ILL | MGI |
|  | INDEL | PacBio | MGI | ILL |
| Difficult-to-map | SNP | PacBio | MGI | 10X |
|  | INDEL | MGI | ILL | PacBio |
| Easy Heterozygous | SNP | PacBio | MGI | ILL |
|  | INDEL | MGI | ILL | PacBio |
| Easy Homozygous Alt | SNP | PacBio | MGI | ILL |
|  | INDEL | ILL | PacBio | MGI |
| Difficult Heterozygous | SNP | PacBio | MGI | ILL |
|  | INDEL | MGI | ILL | PacBio |
| Difficult Homozygous Alt | SNP | PacBio | MGI | ILL |
|  | INDEL | MGI | ILL | PacBio |
| Tier 1 Easy SV | DEL | ONT | PacBio | MGI |
|  | INS | PacBio | ONT | ILL |
| Tier 1+2 Difficult SV | DEL | ONT | PacBio | MGI |
|  | INS | PacBio | ONT | ILL |
| Tier 1 Easy SV GT F1 | DEL | PacBio | ONT | MGI |
|  | INS | PacBio | ONT | MGI |
| Tier 1+2 Diff SV GT F1 | DEL | PacBio | ONT | MGI |
|  | INS | PacBio | ONT | MGI |

### Small Variant Detection

The top-performing technologies for detecting small variants, including SNPs and indels, were SRS platforms such as Illumina and MGI, as well as PacBio among the LRS technologies (**Figure 2**). For easy-to-map SNPs, PacBio, Illumina, and MGI consistently achieved the highest F1-scores, indicating their superior ability to call these variants accurately. Similarly, PacBio and MGI performed well for easy-to-map indels, with Illumina ranking third. This performance was consistent across both hets and homalt variants, although slight differences in ranking were observed depending on the variant type. PacBio was particularly effective at detecting easy-to-map SNPs for both zygosity types, while SRS data excelled in identifying easy-to-map indels with Ilumina ranking highest for alts, and MGI for hets.

In contrast, difficult-to-map regions posed greater challenges, particularly for indels. PacBio, showed the highest F1-scores for difficult-to-map SNPs, indicating that LRS technologies may offer a distinct advantage in these regions due to their longer read lengths and ability to span repetitive or complex genomic sequences. For difficult-to-map indels, however, both SRS platforms outranked PacBio. The performance rankings were consistent for zygosity with the same ordering observed within each variant type category.

Algorithm performance was relatively consistent across SRS platforms especially in easy-to-map regions, with both MGI and ILL achieving similar F1-scores (>99%) for SNPs and indels. However, when focusing on difficult-to-map regions, the sequencing technology choice becomes critical. MGI outperformed Illumina in low-mappability and AT-rich regions, while Illumina was superior in GC-rich regions, and in events around the major histocompatibility complex (MHC) regions. Conversely, algorithm consistency for LRS was more variable especially for ONT, suggesting both technology and algorithm advancements are still required to attain consistent performance as observed in SRS data.

The comparison of small variant calling algorithms across technologies and validation sets (GIAB and CMRG) revealed DeepVariant combined with SRS as the top-performing algorithm by attaining the highest overall F1-score and minimized false discovery rates (FDR), followed by closely by DRAGEN and Clair3. DeepVariant’s precision is critical, as FDR is a key metric in clinical and high-confidence research settings where false positives could lead to misinterpretation of genetic data. DRAGEN, another high-performing variant caller, prioritized sensitivity with the lowest FNR, which makes it preferable when the primary concern is variant discovery, albeit with a slight tradeoff in FDR compared to DeepVariant. Clair3 followed closely and performed well across most technologies, particularly for challenging LRS datasets with PB achieving the highest LRS F1 score. ONT struggled to achieve high accuracy due to its higher susceptibility to sequencing biases, particularly in SNPs and InDels within homopolymer regions. However, ONT calls with Clair3 showed notable performance in SNP detection within difficult-to-map regions. Recent advancements in ONT technology, particularly the introduction of the R10 pore chemistry, have shown significant improvements in mitigating these traditional sources of error. This has led to a marked reduction in indel errors and improved overall basecalling accuracy, which directly translates to better variant calling performance.

Our results suggest that PacBio, while not the top performer overall for indel detection, is particularly effective for SNPs, especially in difficult-to-map regions. This positions PacBio as a versatile technology and underscores the potential utility of combining SRS and LRS technologies for a comprehensive variant detection in both well-mapped and complex regions. Moreover, the recent development of a DeepVariant hybrid model, which integrates both PacBio and SRS data, represents a significant advancement in variant calling. This hybrid model aims to combine the strengths of both SRS and LRS technologies by leveraging PacBio’s high accuracy in detecting difficult-to-map SNPs and complex indels with the higher depth and precision offered by SRS. The hybrid approach showed improved performance for detecting small variants, especially in regions with complex or repetitive sequence structures that typically challenge individual technologies (ref). The ability to incorporate SRS data into this model offers flexibility, as it is expected to be applicable to other SRS platforms like MGI. This emerging hybrid variant calling model is poised to provide a comprehensive framework for improving variant detection accuracy across diverse genomic regions and variant types and merits additional evaluation.

### Structural Variant Detection

To comprehensively assess structural variant (SV) detection and genotyping, we evaluated various sequencing technologies and callers across three validation datasets: genome-wide (Tier 1+2), high-confidence (Tier 1), and clinically relevant (CMRG). LRS technologies consistently outperformed SRS and synthetic sequencing ranking highest in both discovery and genotyping for both variant types. This superior performance can be attributed to the inherent strengths of LRS: ONT excels in resolving large DELs due to its long, continuous reads, while PB, with its high-accuracy reads, is more effective in detecting smaller INS events and achieving higher genotyping fidelity for both variant types. Among LRS callers, Dysgu demonstrated the highest F1 scores in the high-confidence Tier 1 and CMRG datasets, suggesting its suitability for clinical applications. In a research setting, SVIM excelled in discovering and genotyping the largest number of SVs in the more complex Tier 1+2 dataset. In contrast, SRS technologies, while exhibiting higher precision, showed lower recall and overall variant detection accuracy, particularly for INS. Dysgu emerged as the top SRS caller for Tier 1+2, likely due to its ability to identify smaller SVs (30-50 bp range). However, for the high-confidence datasets, the optimal SRS caller varied: Dysgu for INS, DRAGEN for Tier 1 DEL, and MetaSV for CMRG DEL, indicating the importance of caller selection based on dataset characteristics and specific research or clinical goals.

Our analysis of SV size stratification further reinforced the superiority of LRS technologies across all size bins, particularly for smaller SVs of 30-50bp for Tier 1+2 and 50-250bp for Tier 1 and CMGR, where PacBio had the highest accuracy, and larger SVs (>6000 bp) in Tier1+2 and Tier1, where ONT excelled. However, caller performance varied significantly by SV size, validation set, and technology, with Dysgu, SVIM, Sniffles2 and pbsv frequently emerging as top performers for different size categories. SRS technologies, while underperforming overall, showed some strengths in specific size ranges, specifically medium to large deletions (> 500bp). Additionally SRS technologies favored different SV sizes with ILL excelling in small deletions and MGI in larger insertions, however, this could be largely impacted by fragment size selection (insert size) at the library preparation level.

### Optimal Depth for Variant Detection

Coverage depth was a key factor influencing variant detection accuracy across both small variants and SVs. Our analysis showed that a minimum depth of 20x was required for most small variant detection, with more challenging regions and indels benefiting from higher depths. More specially, for SNPs and indels in easy-to-map regions, diminishing returns in variant detection are observed after 15x coverage for PacBio, 20x for SRS, and 30x for ONT. However, in difficult-to-map regions, SRS technologies like Illumina require at least 60x coverage to capture the majority of true variants, particularly for heterozygous calls, which are more challenging to detect than homozygous variants. PacBio HiFi stands out in terms of depth efficiency, achieving >99% sensitivity for both SNPs and InDels at moderate depths (15x–25x) in difficult regions, while SRS and ONT required far greater coverage to reach comparable sensitivity. ONT, despite requiring higher depth (45x–50x), still underperformed, especially for indel detection, where it captured less than 52% of the true positives even at high depths.

Our investigation into the impact of sequencing depth on SV detection revealed that, similar to challenging small variant analysis, increased coverage generally improves SV calling accuracy. However, this improvement exhibits diminishing returns beyond specific thresholds, suggesting that excessively deep sequencing may not always translate to proportional gains in SV detection. Despite the deep sequencing of HG002 samples, all technologies, LRS, still missed a significant number of SVs, particularly in the complex Tier 1+2 and CMRG datasets, highlighting persistent challenges in resolving complex SVs. Within each technology and SV type, optimal performance varied by caller and coverage. For SRS, Dysgu consistently minimized false negatives (FNs) across most SV types and validation sets, typically requiring at least 80x coverage. However, maximizing true positives (TPs) often necessitated even higher coverage, such as 120x for Tier 1 INS. For ONT datasets, a coverage range of 35x to 50x was generally optimal, with SVIM and Sniffles2 demonstrating superior performance for DEL and INS, respectively. Interestingly, for ONT-called INS using SVIM on the Tier 1+2 dataset, lower coverages of 20-25x yielded the highest F1 and recall. PB, while generally benefiting from deeper sequencing (40x to 60x), particularly when using pbsv for INS detection, also showed caller and dataset specific variations. For example, Dysgu required only 15x PB data for optimal INS detection in the CMRG dataset. These findings underscore the importance of optimizing sequencing depth and caller selection based on technology, SV type, and dataset complexity to achieve the most accurate and cost-effective SV detection.

### Clinical and Research Implications

The results of this study have several important implications for clinical genomics and research with the choice of technology, algorithm, and coverage depth should be tailored to the specific genomic context and variant type of interest. For small variant detection in well-characterized regions of the genome, SRS technologies such as Illumina and MGI, coupled with variant callers like DeepVariant or DRAGEN, offer high precision and recall at manageable coverage depths. These technologies are ideal for routine genomic studies and clinical diagnostics, where the detection of well-mapped SNPs and InDels is a primary concern. However, for more challenging regions and SVs, LRS technologies like PacBio and ONT are indispensable, but under certain conditions such as higher coverage and a priori knowledge of the variants of interest (for example SVs of size larger than 500bp) SRS can be applicable. However, LRS technologies’ ability to span both small and large deletions and insertions, combined with high genotype accuracy, makes them essential for capturing the full spectrum of human genetic variation, especially in clinical applications where SVs are implicated in disease.

## Conclusion

This study provides a comprehensive evaluation of sequencing technologies and variant calling algorithms for accurately detecting SNPs, InDels, and SVs. Our findings highlight the strengths and limitations of different approaches, informing optimal technology and methodology choices for various genomic contexts and research objectives. While SRS technologies excel in detecting small variants in well-mapped regions, LRS technologies, particularly PacBio, offer superior performance for SVs and difficult-to-map regions.

Future advancements in sequencing technology, such as continued improvements in ONT and Pacbio chemistries (R10.4.1 and SPRQ) and the development of more accurate indel calling algorithms, are expected to further enhance the performance of LRS platforms. Additionally, the transition to pangenome alignments, which incorporate reference variation from diverse populations, holds the promise of improving variant detection accuracy, particularly in underrepresented populations. By leveraging these advancements, researchers will be better equipped to uncover the full spectrum of human genetic variation and gain valuable insights into disease mechanisms, population genetics, and evolutionary biology.

## Data availability

The data used to support the findings of this study are available from the corresponding author upon request.

## Materials and Methods

### Library preparation and sequencing

A large batch of cells expanded from one original vial directly obtained from the Coriell Institute for Medical Research cell line repository was used for DNA isolation and subsequent genomic analysis using four different technology platforms: Illumina HiSeq Xten (ILL), MGI DNBSEQ-400 (MGI), Oxford Nanopore Technologies (ONT1), and 10x genomics (sequenced on Illumina HiSeq Xten; 10X).

Additional LRS data was downloaded directly from Genome in a Bottle (GIAB) for additional analyses. Two Pacific Biosciences (PB) HiFi datasets: PacBio CCS 15kb (PB1) and PacBio CCS 15kb and 20kb chemistry 2 (PB2) and one Oxford Nanopore Technologies (ONT2)

### Cell line culture

GM24385 (H0002), GM24149 (HG003), and GM24143 (HG004) cells (Coriell Institute) were cultured in RPMI medium supplemented with 2 mM L-alanyl-L-glutamine, 15% fetal bovine serum, 50 U/ml penicillin, and 50 µg/ml streptomycin (all from ThermoFisher Scientific, Nepean, ON, Canada) at 37°C and 5% CO_2_. Cells were seeded at a density of 500,000 cells/ml and passaged or harvested when they reached 1 M cells/ml.

### DNA extraction

Cells were extracted on the SageHLS High Molecular Weight DNA Library System (Sage Science, Inc., Beverly, Massachusetts, United States) which aims to minimize pipetting steps that shear very HMW DNA by doing extraction and fractionation of DNA based in the same cassette. Cells in suspension are lysed directly in the cassette and an enzyme is added to digest the DNA just enough so that it can be electrophoresed and separated by size. This digestion step is crucial, as overshearing will greatly reduce the yield of HMW fragments. A second electrophoresis is then done perpendicular to the first, in order to collect the DNA in 6 different elution wells.

#### PCRfree-Whole Genome Sequencing Library Preparation

##### Illumina

The Ashkenazi trio genomic DNA (gDNA) quality was assessed prior to library preparation to ensure high-quality input material. DNA integrity was evaluated using either agarose gel electrophoresis or the Agilent Tapestation, while quantification was performed using the Qubit DNA High Sensitivity (HS) assay. DNA purity was assessed by measuring the A260/280 ratio, with acceptable values ranging between 1.8 and 2.0. Whole-genome sequencing (WGS) libraries were prepared using the Illumina TruSeq PCR-Free Library Prep Kit, with 500 ng of high-quality gDNA as input. DNA was fragmented to a target size of 300 bp using the Covaris LE220 focused ultrasonicator, followed by size selection with Ampure XP beads (Beckman Coulter). Library quality control (QC) was performed to ensure appropriate fragment size and concentration before sequencing. Fragment size distribution was assessed using the Agilent Bioanalyzer DNA High Sensitivity assay, while library quantification was performed using the KAPA qPCR Library Quantification Kit. Sequencing was carried out on the Illumina HiSeq X platform with paired-end 150 bp reads (2 × 150), on one lane with 1% PhiX as an internal control.

##### MGI

The samples were done using the WGS Lucigen (NxSeq AmpFREE low DNA Library Kit) with the starting input of 1000ng .The Genomic DNA was mechanically sheared using COVARIS at 300bp. Libraries were quantified using the KAPA Library Quant Kit (Illumina/Universal qPCR) from Roche for qPCR quantification. Average size fragment was determined using a LabChip GXII (PerkinElmer) instrument. The libraries were converted to MGI using the MGIEasy Universal Library Conversion Kit (App-A) using 50 ng of each sample with 5-cycles of amplification. The converted libraries were quantified using Qubit™ ssDNA Assay Kit and pooled at equimolar. The pool was loaded at 60 fmol on a DNBSEQ-G400 as per the manufacturer’s recommendations. The run was performed for 2x150 cycles (paired-end mode).

#### 10X Genomics

The DNA extractions for HG002 and HG004 were prepared following the SageHLS High Molecular Weight DNA Extraction protocol using a 1/800 dilution of NEBNext® dsDNA Fragmentase® (cat# M0348) for a half hour digestion. DNA from HG003 was prep testing an alternative protocol which uses a Cas9 system (Cas9 Nuclease, *S. pyogenes*, New England BioLabs, Inc., cat# M0386) with two simple guide RNAs (a 10 CA repeat guide and a 10 AT repeat guide, Alt-R® CRISPR-Cas9 crRNA, Integrated DNA Technologies, Inc., Coralville, Iowa, United States). DNA yield from each elution well was measured by Qubit™ dsDNA BR Assay Kit (ThermoFisher Scientific, cat# Q32853) and molecule length was assessed by Femto Pulse (Genomic DNA 165 kb Kit, 3 hour run, Agilent Technologies, Inc., Santa Clara, California, United States, cat# FP-1002-0275).

#### Nanopore

Genomic DNA was manually purified from whole blood using the Qiagen protocol for high-molecular-weight (HMW) DNA (Qiagen, Toronto, ON, Canada). Briefly, cells were pelleted by centrifuging at 500×g for 10 minutes at 4°C, and the supernatant was removed, leaving approximately 200 µL with the pellet. The pellet was then resuspended in a digestion mixture containing 20 µL proteinase K, 4 µL RNase A, and 150 µL Buffer AL, vortexed, and incubated at room temperature for 30 minutes. Following a quick spin, DNA was bound to 15 µL MagAttract Suspension G by adding 280 µL Buffer MB and incubating for 3 minutes at 1,400 rpm at room temperature in a thermoshaker. After a quick spin, the supernatant was removed after a minute incubation on a magnetic rack. The magnetic beads were then washed twice with 700 µL Buffer MW1 and twice with 700 µL Buffer PE, each wash involving a 1-minute incubation at 1,400 rpm in the thermoshaker followed by 1 minute on the magnetic rack and supernatant removal. The beads were further rinsed twice with 700 µL of water while on the magnetic rack, removing the supernatant after 1 minute each time. Finally, the DNA was eluted by adding 100 µL Buffer AE to the beads (removed from the magnetic rack), incubating at room temperature for 3 minutes at 1,400 rpm, placing tubes on the magnetic rack, and recovering the supernatant into a new tube after 1 minute. Purified DNA was stored at 4°C, and its quality was assessed using a NanoDrop 8000 Spectrophotometer (ThermoFisher Scientific, Nepean, ON, Canada) The Qiagen protocol for manual purification of high-molecular-weight (HMW) genomic DNA from whole blood was used as recommended by the manufacturer (Qiagen, Toronto, ON, Canada).

ONT1 library preparation was performed using 1.5 µg genomic DNA as starting material for the ligation sequencing kit (SQK-LSK109) as recommended by the manufacturer (Oxford Nanopore Technologies, Oxford, UK). The library was loaded onto a FLO-PRO002 R9.4.1 flow cell with 6052 available pores at the start of the sequencing run. Flow cell was refueled with 150 µl priming mix after ∼45 h of sequencing. PromethIONBeta Release 19.05.1 was used with live high accuracy basecalling (Guppy v3.0.3).

Sequencing was primarily performed on the MinION using a mix of SQK-RAD003 and SQK-RAD004 library prep kits on the FLO-MIN106. Additional sequencing was performed using the GridION and PromethION. Reads were base-called with guppy (version 2.3.5), then combined into a single fastq file. The high accuracy basecalling algorithm was used for all but one flowcells (FAH03763), as the model config file was not available for that particular flowcell/ sequencing kit combination (flowcell - FLO-MIN107 and kit - SQK-RAD003).

The ONT2 “ultralong" (Jain, et al 2018) libraries were generated on GIAB Ashkenazi son cell line HG002 (GM24385) to generate a library with 52-fold total coverage with approximately 15-fold coverage generated by reads larger than 100kb. DNA was prepared using various modified versions of Josh Quick’s protocol (https://dx.doi.org/10.17504/protocols.io.mrxc57n). A manuscript with a detailed protocol and methods development is in prep. Sequencing was primarily performed on the MinION using a mix of SQK-RAD003 and SQK-RAD004 library prep kits on the FLO-MIN106 with additional sequencing performed on GridION and PromethION. Reads were base-called with guppy (version 3.4.5), then combined into a single fastq file. The hac algorithm was used for all but one flowcells (FAH03763), as the model config file was not available for that particular flowcell/ sequencing kit combination (flowcell - FLO-MIN107 and kit - SQK-RAD003)

#### PacBio

High-fidelity 15kb long read dataset of HG002 (PB1) was generated by the GIAB consortium using using Binding/Sequencing chemistry 3.0 and SMRT Link 6.0. One shotgun library was prepared from a DNA sample extracted from a large homogenized growth of B-lymphoblastoid cell lines from the Coriell Institute for Medical Research. Size selection was performed, targeting a narrow size band ∼15kb using the sageELF DNA size-selection system from SAGE Science. Circular consensus sequences were calculated using default parameters and a predicted read accuracy filter of Q20 (99%) using SMRT Link 6.0. The library was sequenced to approximately 28-fold coverage.

High-fidelity combined 15kb and 20kb long read dataset of HG002 (PB2) was generated by the GIAB consortium using using Sequel II System with 2.0 chemistry and SMRT Link 9.0. Two shotgun library were prepared from a DNA sample extracted from a large homogenized growth of B-lymphoblastoid cell lines from the Coriell Institute for Medical Research the first with TPK library prep and SageELF Fraction 4 (target: 15 kb) size selection and the second with SMRTbell Express 2.0 + Enzyme Clean Up and SageELF Fraction 2 (target: 20 kb) size selection. Circular consensus sequences were calculated using default parameters and a predicted read accuracy filter of Q20 (99%) using SMRT Link 9.0. The 15kb library consisting of four readsets was sequenced to approximately 36-fold coverage and the 20kb library with two readset totaling 16-fold coverage.

### Bioinformatics Methods

#### Experimental Design

This study aimed to evaluate the performance of various sequencing technologies and variant calling algorithms in detecting germline SNPs, InDels, and SVs in human genomes. SRS platforms, including Illumina and MGI, LRS technologies such as PacBio and ONT, and synthetic 10X genomics, were used to generate sequencing data. All reads were linearly aligned to the human genome build GRCh37 decoy (hs37d5) to ensure consistency across datasets. A range of variant callers were applied, some optimized for specific sequencing technologies, to assess their ability to accurately detect variants. Downsampling experiments were conducted to further evaluate the effects of sequencing depth on variant detection.

## Data Processing

### SRS Data Processing

Short-read sequencing data from Illumina and MGI platforms were processed using GenPipes version 4.5.0, a bioinformatics pipeline based on GATK Best Practices. Quality control of raw reads was performed using Skewer (v0.2.2), followed by alignment to the GRCh37 decoy (hs37d5) reference genome using the BWA-MEM algorithm (v7.15). Post-alignment processing included marking duplicate reads using Picard (v2.9.0) and performing base quality score recalibration with GATK (v3.8). Parallel alignments were generated for comparison using Parabrick (v2.4.0-rc3), DRAGEN (v07.011.350.3.3.11), and MegaBolt (v2.2.2.1), with each tool run using default settings.

### LRS Data Processing

Long-read sequencing data generated from PacBio and ONT platforms were aligned to the GRCh37 decoy (hs37d5) genome using technology-specific aligners. ONT reads were aligned using Minimap2 (v2.26) with parameters optimized for contiguity (-a -z 600,200 -x map-ont), while PacBio long reads were aligned using pbmm2 (v1.13.1) with the circular consensus sequencing (CCS) preset. Datasets generated both in-house and from public repositories were processed with the same protocols to ensure consistency across samples.

## Downsampling

To evaluate the effects of sequencing depth on variant detection, downsampling experiments were performed on both SRS and LRS datasets. Illumina data for the Ashkenazim trio (GM24385/HG002, HG003, HG004) were combined to achieve mean coverage levels of 126x, 120x, and 111x, respectively, and then downsampled to 11 individual coverage levels (ranging from 120x to 2x) using a custom Perl script to randomly select read pairs. The MGI HG002 sample was similarly downsampled into nine coverage levels (from 40x to 2x). For LRS datasets, PacBio (PB2, 62x) and ONT (ONT2, 52x) data were downsampled into 12 and 11 samples, respectively, using rasusa (v2.1.0) which accounted for read-length as part of its downsampling calculation.

## Data Analysis

### Germline Variant Calling

Germline variants, including SNPs and InDels, were called across the SRS datasets using seven variant calling algorithms: Clair3 (v1.0.4), DeepVariant (v1.2.0), DRAGEN, GATK HaplotypeCaller (v3.8 and v4.1.2.0), Parabrick, and MegaBolt. These tools were run with default settings to ensure a consistent comparison across platforms. For the LRS datasets, germline variants were identified using Clair3, DeepVariant, and GATK (v4.1.2.0) for PacBio data, and Clair3, DeepVariant, Pepper-Margin-DeepVariant, and Nanocaller (v2.0.0) for ONT data. All variant callers were executed with default parameters unless otherwise specified.

### Structural Variant Detection

SVs were identified using distinct workflows for SRS and LRS data. For the SRS datasets, structural variants were detected using six SV callers—delly, lumpy, manta, WHAM, breakseq2, and cnvkit—integrated through the MetaSV (v0.5.4) ensemble approach. MetaSV was configured to enhance insertion detection with the --boost_ins option and improve breakpoint resolution. Further filtering was applied using DupHold (v0.2.1), with a fold-change threshold (DHFFC) of 0.7 to minimize false positives for deletions and duplications. Additionally, DRAGEN’s SV pipeline which used Manta and Dysgu (v1.6.1) was applied to the SRS datasets. For both PacBio and ONT LRS datasets, five SV callers—CuteSV (v2.0.3), Dysgu, pbsv (for PacBio-only; 2.9.0), Sniffles (v2.2.2), and SVIM (v2.0.0)—were employed with a minimum SV size of 30 base pairs.

### Germline Variant Assessment

The precision, sensitivity, and F1-scores of germline SNP and InDel calls were evaluated using the hap.py (v0.3.12) algorithm and RTG vcfeval (v3.6.10). Variant calls from each sequencing technology and caller were compared against the Genome in a Bottle (GIAB) v4.2.1 and Clinically Relevant Molecular Genetics (CMRG) v1.0.0 truth sets. This analysis provided a comprehensive assessment of variant detection accuracy across technologies, including comparisons of individual variant callers.

### Structural Variant Benchmarking

The performance of SV detection was benchmarked using the SVbench (v1.0.0) Python API, based on comparisons to three curated truth sets: Tier 1+2 (complex SVs) and Tier 1 (easy SVs) from the GIAB SV truth set (v0.6) and the CMRG subset of clinical SVs. The Tier 1+2 truth set included 15,001 deletions and 15,948 insertions, all over 30 bp in length. The Tier 1 subset contained a smaller number of sequence-resolved SVs, with 4,117 deletions and 5,261 insertions over 50 bp. The CMRG subset encompassed 89 high-quality deletions and 105 insertions located near 273 known clinical genes. Benchmarking metrics, including precision, recall, and F1-scores, were generated for each sequencing technology, and SV size stratification was applied. SVs were categorized into the following size ranges: 30-49 bp, 50-250 bp, 251-500 bp, 501-6000 bp, and 6001-260000000 bp, based on known repeat sizes from GIAB.

### Genotype Performance

Genotype accuracy was assessed by calculating genotype F1-scores for each caller and technology. Additional stratification of results based on SV size allowed for detailed performance comparisons, enabling insight into the genotype calling ability across various types and sizes of SVs.

## Acknowledgements

This work was supported by a Canada Institute of Health Research (CIHR) project grant (PJT-191707). G.B. is supported by a Canada Research Chair Tier 1 award and a FRQ-S Distinguished Research Scholar award. The Canadian Center for Computational Genomics (C3G) was supported by a Genome Canada Genome Technology Platform grant. This research was also enabled in part by support provided by Calcul Québec and the Digital Research Alliance of Canada.

We would like to thank Dr. Albena Pramatarova for providing the cells, and we would like to thank Dr. Pierre Berube for the library preparation, and Shu-Huang Chen and Julio Serna Vasquez for excellent technical assistance.

