## Supplemental_figures_and_tables for "Finding an optimal sequencing strategy to detect short and long genetic variants in a human genome"

### *Supplementary figures*

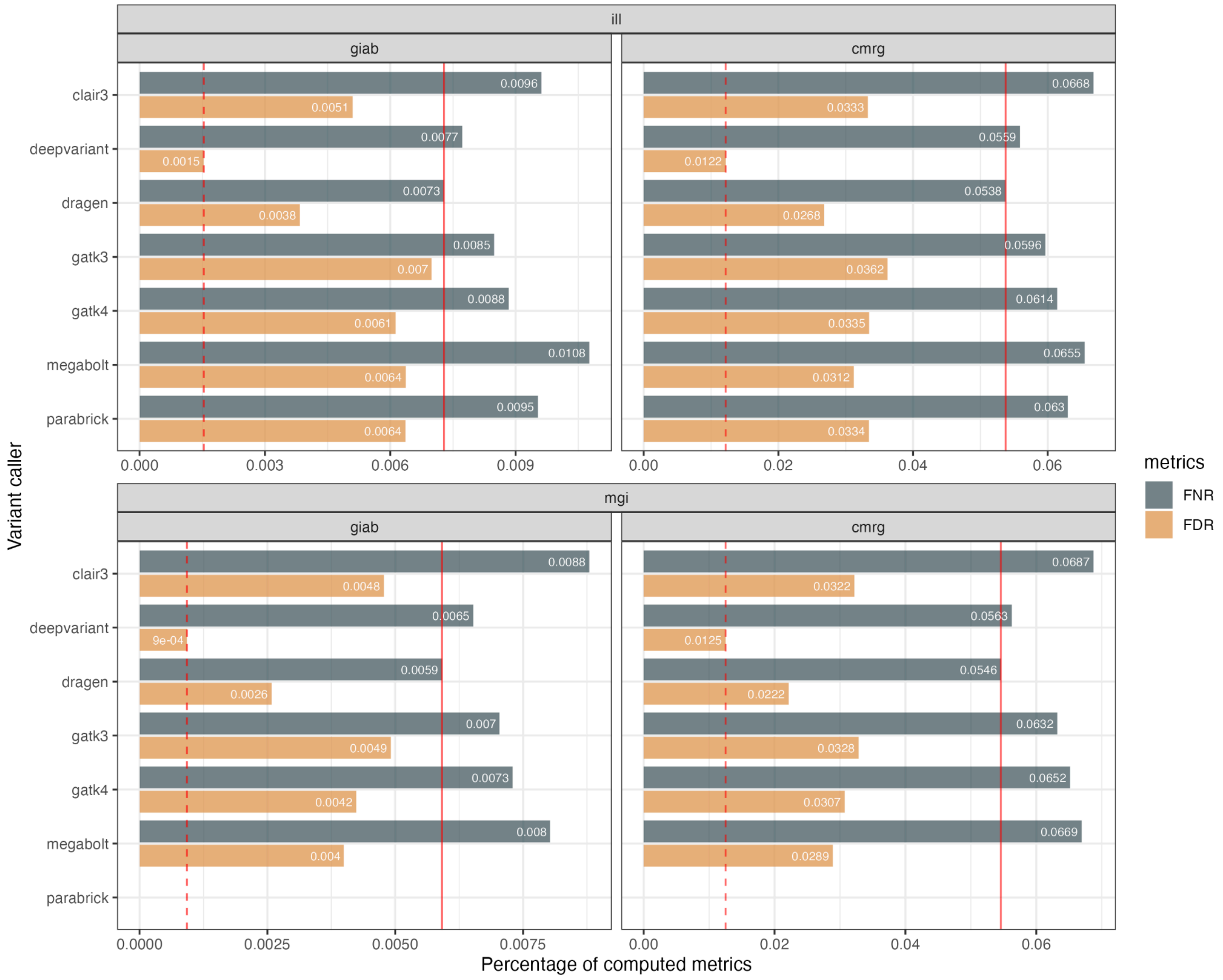


***Figure S1***

Comparing the FNR and FDR for both GIAB (4.2.1) and CMGR validation sets across diploid germline callers using one lane of 41x Illumina Xten (ILL) and 44x MGI G400 sequencing data. Each label represents either the FN or FD rates with a dash or solid vertical line to represent to minimal values representing the top ranked performer

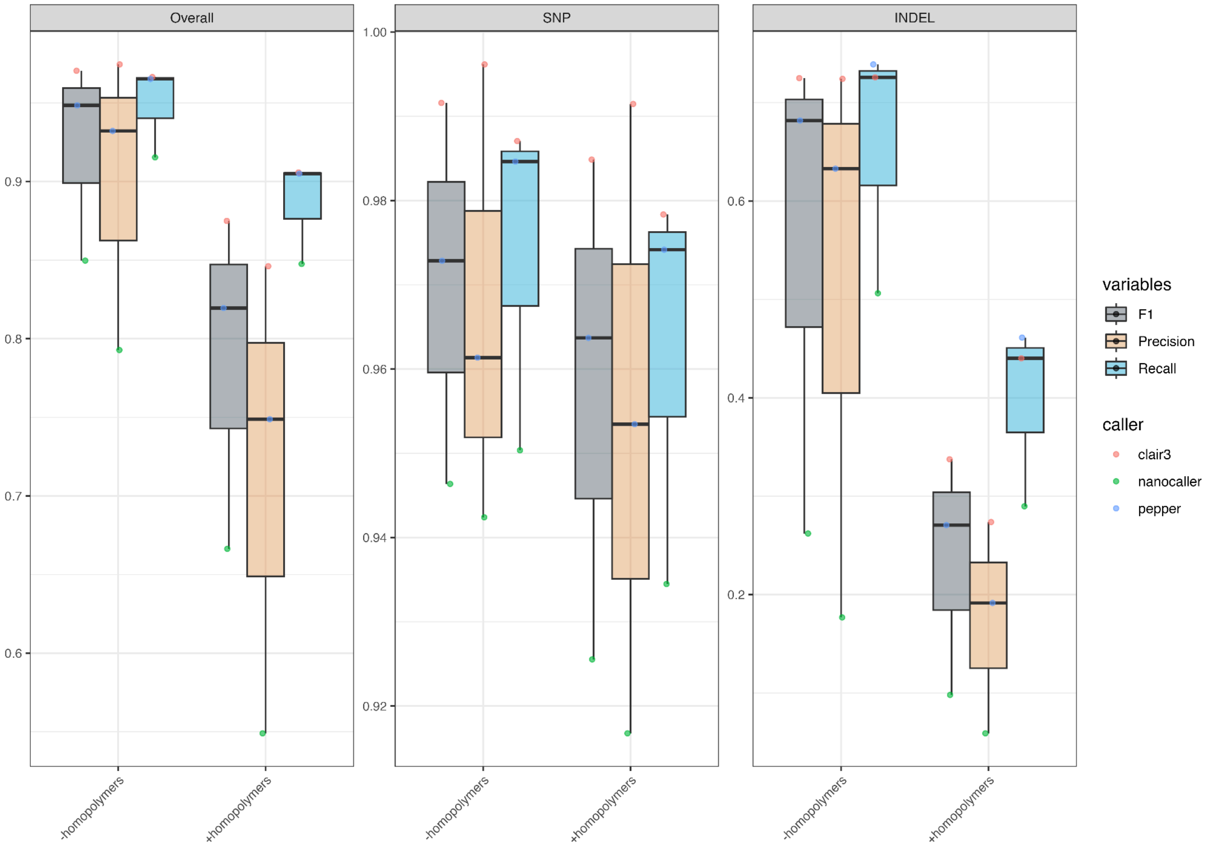


#### *Figure S2*

A boxplot graph comparing three small variant callers: Clair3, Nanocaller and Pepper-Margin-DeepVariant (pepper) for F1, Precision and Recall metrics using the R9.4 ONT1 sample in the presence and absence of variant within proximity homopolymer regions; a known sequence bias in ONT data which drastic affects the performance of variant calling in InDels particularly.

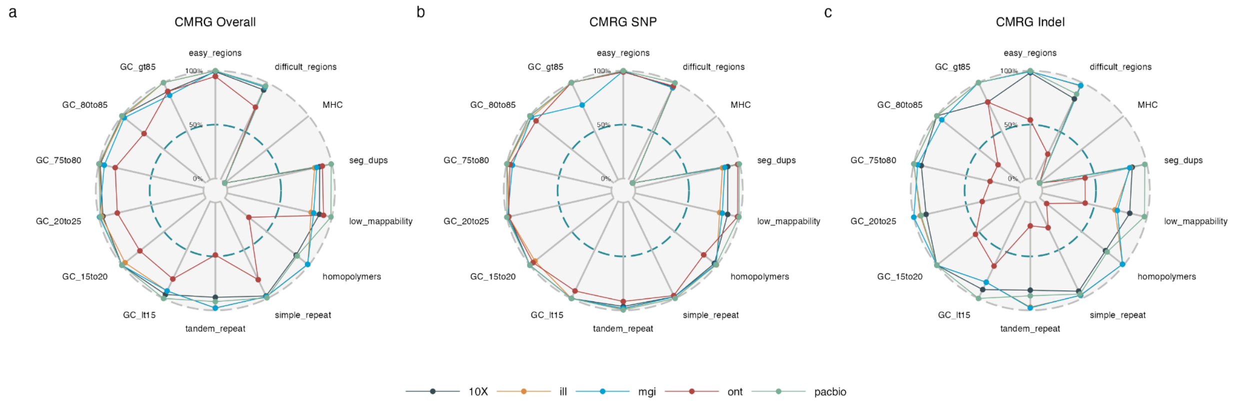

#### *Figure S3*

F1 scores across all tested technologies: SRS (MGI and Illumina), LRS (PacBio and ONT) and synthetic (10X) stratified by GA4GH easy and difficult to map regions for a) GIAB dataset for both SNPs and InDels combined, b) GIAB SNPs only c) GIAB InDels only d) CMRG dataset for both SNPs and InDels combined, e) CMRG SNPs only and f) CMRG InDels only

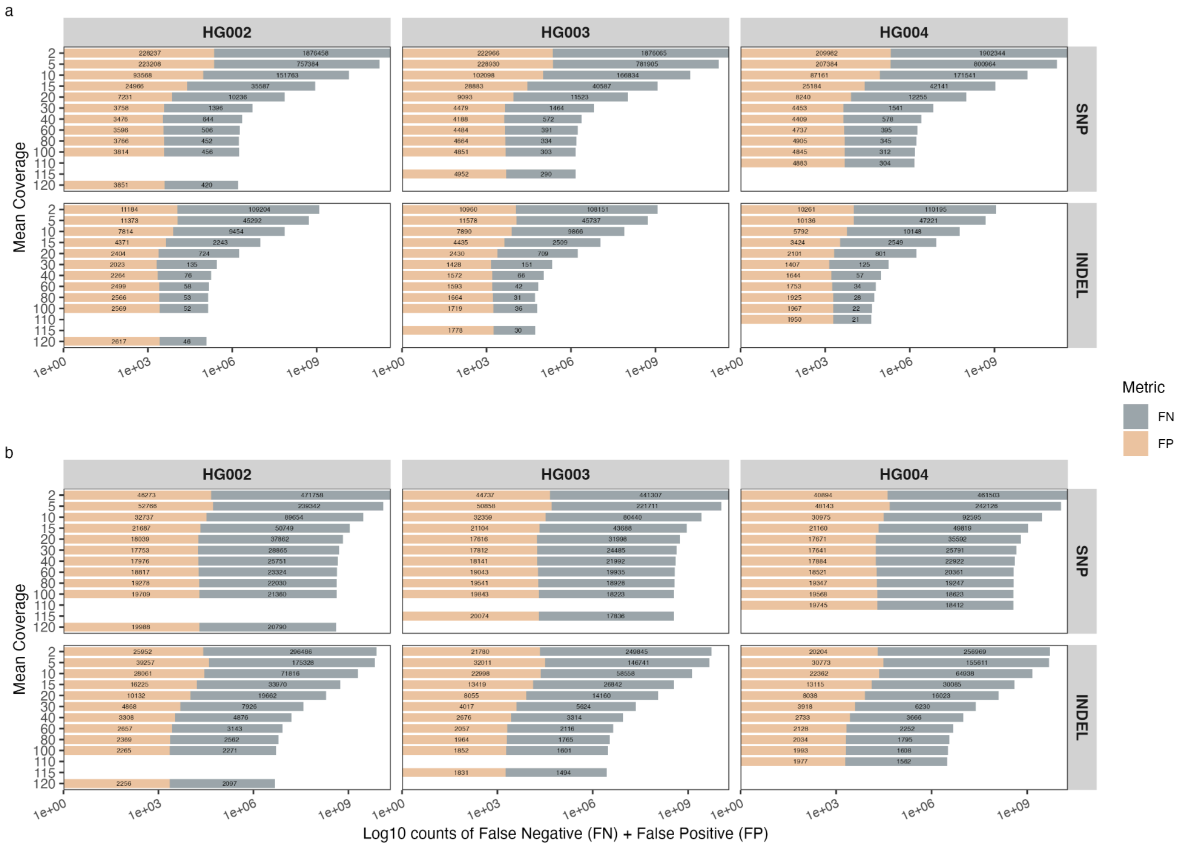


#### *Figure S4*

Downsampling of three lanes HiSeq X sequencing for each sample (HG002/GM24385, HG003/GM24143, and HG004/GM24149) of the Ashkenazim Trio from 2- to 120-fold coverage. False positive and False negative counts were collected for the a) easy-to-map regions. b) difficult-to-map regions.

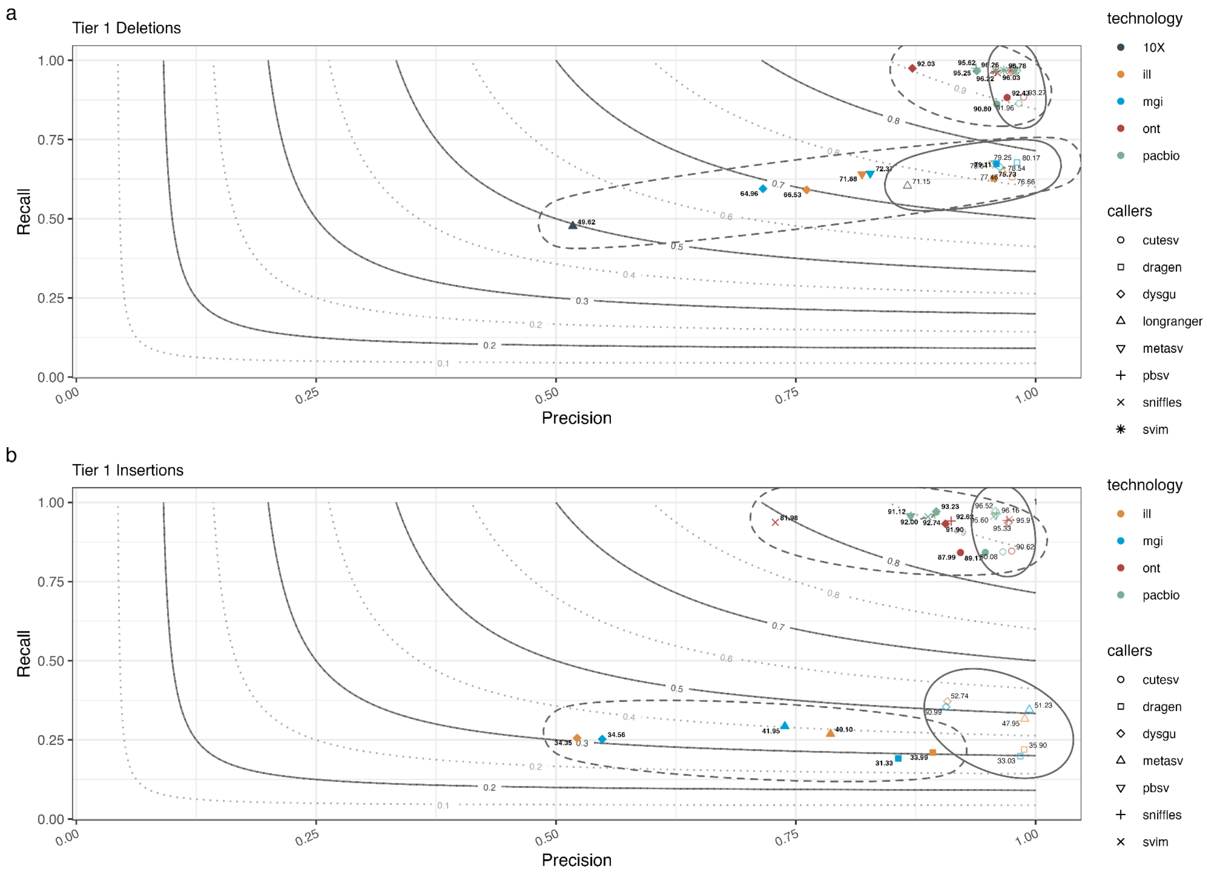


***Figure S5***

Precision-recall curves with discovery F1 (solid line ellipses) and genotyping F1 (dashed line ellipses) across the high quality Tier 1 validation set. Tier 1 contain 4117 deletions a) and 5261 insertions b).

Tier 1 contain 4117 and 5261 high quality deletions b) and insertions d),

#### ***Supplementary Tables***

| | | Depth of Coverage | | | Insert length distribution | | |
|---|---|---|---|---|---|---|---|
| technology | instrument | median | mean | sd | median | mean | sd |
| Illumina | Xten | 42 | 40.9 | 12.7 | 293 | 308.8 | 117.2 |
| MGI | G400 | 45 | 44.6 | 12.1 | 371 | 378.3 | 84.8 |
| Nanopore | PromethION | 29 | 29.2 | 7.8 | 4000 | 7177.8 | 10531.9 |
| PacBio | HiFi | 29 | 29.1 | 8.4 | 13000 | 12971.4 | 1222.4 |
| 10X Genomics | | 26 | 27.3 | 53.2 | 34000 | 45884.7 | 41749.8 |

***Table S1***
Coverage, insert or read size metrics for all HG002 samples used in this study

| | GIAB version 4.2.1 | | | CMRG version 1.0.0 | | |
|---|---|---|---|---|---|---|
| caller | Precision | Recall | F1 score | Precision | Recall | F1 score |
| ***Combined*** | | | | | | |
| **Illumina** | | | | | | |
| Clair3 | 99.49% | 99.04% | 99.26% | 96.67% | 93.20% | 94.90% |
| deepvariant | **99.85**% | 99.23% | **99.54**% | **98.78**% | 94.28% | **96.48**% |
| DRAGEN | 99.62% | **99.27**% | 99.44% | 97.32% | **94.50**% | 95.89% |
| GATK3 | 99.30% | 99.15% | 99.23% | 96.38% | 93.94% | 95.14% |
| GATK4 | 99.39% | 99.12% | 99.25% | 96.65% | 93.76% | 95.18% |
| megabolt | 99.36% | 98.92% | 99.14% | 96.88% | 93.35% | 95.08% |
| parabrick | 99.36% | 99.05% | 99.21% | 96.66% | 93.60% | 95.10% |
| **MGI** | | | | | | |
| Clair3 | 99.52% | 99.12% | 99.32% | 96.78% | 93.00% | 94.85% |
| deepvariant | **99.91**% | 99.35% | **99.63**% | **98.75**% | 94.25% | **96.45**% |
| DRAGEN | 99.74% | **99.41**% | 99.57% | 97.78% | **94.44**% | 96.08% |
| GATK3 | 99.51% | 99.30% | 99.40% | 96.72% | 93.58% | 95.12% |
| GATK4 | 99.58% | 99.27% | 99.42% | 96.93% | 93.38% | 95.12% |
| megabolt | 99.60% | 99.20% | 99.40% | 97.11% | 93.21% | 95.12% |
| ***SNP*** | | | | | | |
| **Illumina** | | | | | | |
| Clair3 | 99.63% | 99.20% | 99.41% | 97.31% | 94.06% | 95.66% |
| deepvariant | **99.87**% | 99.25% | **99.56**% | **99.01**% | 94.59% | **96.75**% |
| DRAGEN | 99.66% | **99.30**% | 99.48% | 97.55% | **94.75**% | 96.13% |
| GATK3 | 99.35% | 99.19% | 99.27% | 96.39% | 94.23% | 95.30% |
| GATK4 | 99.43% | 99.16% | 99.29% | 96.71% | 94.11% | 95.39% |
| megabolt | 99.50% | 99.09% | 99.29% | 97.08% | 93.97% | 95.50% |
| parabrick | 99.45% | 99.14% | 99.30% | 96.76% | 94.06% | 95.39% |
| **MGI** | | | | | | |
| Clair3 | 99.63% | 99.25% | 99.43% | 97.44% | 93.91% | 95.65% |
| deepvariant | **99.92**% | 99.34% | **99.63**% | **98.96**% | 94.74% | **96.81**% |

| | | | | | | |
|---|---|---|---|---|---|---|
| DRAGEN | 99.75% | **99.40**% | 99.58% | 98.02% | **94.81**% | 96.39% |
| GATK3 | 99.48% | 99.30% | 99.39% | 96.72% | 94.01% | 95.35% |
| GATK4 | 99.56% | 99.27% | 99.42% | 97.00% | 93.85% | 95.40% |
| megabolt | 99.62% | 99.23% | 99.42% | 97.17% | 93.80% | 95.45% |
| ***Indels*** | | | | | | |
| **Illumina** | | | | | | |
| Clair3 | 98.57% | 98.02% | 98.29% | 93.49% | 88.98% | 91.18% |
| deepvariant | **99.67**% | **99.07**% | **99.37**% | **97.65**% | 92.80% | **95.16**% |
| DRAGEN | 99.31% | **99.07**% | 99.19% | 96.17% | **93.26**% | 94.69% |
| GATK3 | 99.00% | 98.92% | 98.96% | 96.33% | 92.49% | 94.37% |
| GATK4 | 99.11% | 98.86% | 98.99% | 96.37% | 92.06% | 94.16% |
| megabolt | 98.47% | 97.88% | 98.17% | 95.89% | 90.34% | 93.03% |
| parabrick | 98.81% | 98.42% | 98.62% | 96.12% | 91.35% | 93.67% |
| **MGI** | | | | | | |
| Clair3 | 98.86% | 98.32% | 98.59% | 93.49% | 88.54% | 90.95% |
| deepvariant | **99.84**% | 99.39% | **99.62**% | **97.68**% | 91.84% | **94.67**% |
| DRAGEN | 99.69% | **99.46**% | 99.57% | 96.61% | **92.66**% | 94.59% |
| GATK3 | 99.66% | 99.30% | 99.48% | 96.68% | 91.46% | 94.00% |
| GATK4 | 99.67% | 99.26% | 99.47% | 96.58% | 91.10% | 93.76% |
| megabolt | 99.50% | 98.96% | 99.23% | 96.81% | 90.34% | 93.46% |

***Table S2***

Illumina and MGI data. Results were calculated based on the GIAB and CMGR validation datasets.

| | | Processing walltime | | |
|---|---|---|---|---|
| Technology | Caller | bam processing | variant calling | total time |
| **Illumina** | | | | |
| | GATK3 | 30:43:46 | 5:01:58 | 35:45:44 |
| | GATK4 | 30:43:46 | 2:40:35 | 33:24:21 |
| | DeepVariant | 30:43:46 | 1:01:38 | 31:45:24 |
| | Dragen | 0:19:54 | 0:13:18 | 0:33:12 |
| | MegaBolt | 3:07:39 | 1:36:38 | 4:44:17 |
| | Parabrick | 0:56:50 | 0:20:03 | 1:16:53 |
| **MGI** | | | | |

| | | | | |
|---|---|---|---|---|
| | GATK3 | 41:57:11 | 4:32:00 | 46:29:11 |
| | GATK4 | 41:57:11 | 7:05:47 | 49:02:58 |
| | DeepVariant | 41:57:11 | 0:52:58 | 42:50:09 |
| | Dragen | 0:20:16 | 0:13:41 | 0:33:57 |
| | MegaBolt | 2:11:54 | 5:28:38 | 7:40:32 |

***Table S3***
Computational resources.

| | GIAB version 4.2.1 | | | CMRG version 1.0.0 | | |
|---|---|---|---|---|---|---|
| caller | Precision | Recall | F1 score | Precision | Recall | F1 score |
| ***Combined*** | | | | | | |
| **Illumina** | | | | | | |
| clair3 | 99.49% | 99.04% | 99.26% | 96.67% | 93.20% | 94.90% |
| deepvariant | 99.85% | 99.23% | 99.54% | 98.78% | 94.28% | 96.48% |
| dragen | 99.62% | 99.27% | 99.44% | 97.32% | 94.50% | 95.89% |
| gatk4 | 99.39% | 99.12% | 99.25% | 96.65% | 93.76% | 95.18% |
| **MGI** | | | | | | |
| clair3 | 99.52% | 99.12% | 99.32% | 96.78% | 93.00% | 94.85% |
| deepvariant | 99.91% | 99.35% | 99.63% | 98.75% | 94.25% | 96.45% |
| dragen | 99.74% | 99.41% | 99.57% | 97.78% | 94.44% | 96.08% |
| gatk4 | 99.58% | 99.27% | 99.42% | 96.93% | 93.38% | 95.12% |
| **10X** | | | | | | |
| longranger | 98.73% | 96.92% | 97.82% | 96.69% | 89.39% | 92.90% |
| **Nanopore** | | | | | | |
| clair3 | 84.61% | 90.58% | 87.49% | 84.75% | 85.37% | 85.06% |
| nanocaller | 54.90% | 84.76% | 66.64% | 50.59% | 78.24% | 61.45% |
| pepper | 74.88% | 90.50% | 81.95% | 74.29% | 84.89% | 79.23% |
| **Pacbio** | | | | | | |
| clair3 | 98.32% | 99.16% | 98.74% | 96.55% | 96.69% | 96.62% |
| deepvariant | 97.38% | 98.76% | 98.06% | 95.43% | 95.97% | 95.70% |
| gatk4 | 78.96% | 97.54% | 87.27% | 78.89% | 93.75% | 85.68% |
| ***SNP*** | | | | | | |
| **Illumina** | | | | | | |
| clair3 | 99.63% | 99.20% | 99.41% | 97.31% | 94.06% | 95.66% |

| | | | | | | |
|---|---|---|---|---|---|---|
| deepvariant | 99.87% | 99.25% | 99.56% | 99.01% | 94.59% | 96.75% |
| dragen | 99.66% | 99.30% | 99.48% | 97.55% | 94.75% | 96.13% |
| gatk4 | 99.43% | 99.16% | 99.29% | 96.71% | 94.11% | 95.39% |
| **MGI** | | | | | | |
| clair3 | 99.63% | 99.25% | 99.43% | 97.44% | 93.91% | 95.65% |
| deepvariant | 99.92% | 99.34% | 99.63% | 98.96% | 94.74% | 96.81% |
| dragen | 99.75% | 99.40% | 99.58% | 98.02% | 94.81% | 96.39% |
| gatk4 | 99.56% | 99.27% | 99.42% | 97.00% | 93.85% | 95.40% |
| **10X** | | | | | | |
| longranger | 99.65% | 98.78% | 99.21% | 98.39% | 93.30% | 95.78% |
| **Nanopore** | | | | | | |
| clair3 | 99.15% | 97.83% | 98.49% | 98.09% | 95.35% | 96.70% |
| nanocaller | 91.68% | 93.45% | 92.55% | 84.79% | 89.94% | 87.29% |
| pepper | 95.35% | 97.42% | 96.37% | 92.96% | 94.56% | 93.75% |
| **Pacbio** | | | | | | |
| clair3 | 99.93% | 99.91% | 99.92% | 99.31% | 98.65% | 98.98% |
| deepvariant | 99.94% | 99.88% | 99.91% | 99.46% | 98.52% | 98.99% |
| gatk4 | 99.39% | 99.52% | 99.46% | 98.83% | 96.94% | 97.88% |
| ***INDELS*** | | | | | | |
| **Illumina** | | | | | | |
| clair3 | 98.57% | 98.02% | 98.29% | 93.49% | 88.98% | 91.18% |
| deepvariant | 99.67% | 99.07% | 99.37% | 97.65% | 92.80% | 95.16% |
| dragen | 99.31% | 99.07% | 99.19% | 96.17% | 93.26% | 94.69% |
| gatk4 | 99.11% | 98.86% | 98.99% | 96.37% | 92.06% | 94.16% |
| **MGI** | | | | | | |
| clair3 | 98.86% | 98.32% | 98.59% | 93.49% | 88.54% | 90.95% |
| deepvariant | 99.84% | 99.39% | 99.62% | 97.68% | 91.84% | 94.67% |
| dragen | 99.69% | 99.46% | 99.57% | 96.61% | 92.66% | 94.59% |
| gatk4 | 99.67% | 99.26% | 99.47% | 96.58% | 91.10% | 93.76% |
| **10X** | | | | | | |
| longranger | 92.35% | 84.99% | 88.52% | 86.91% | 70.26% | 77.70% |
| **Nanopore** | | | | | | |
| clair3 | 27.37% | 44.01% | 33.75% | 30.98% | 36.56% | 33.54% |
| nanocaller | 5.90% | 28.97% | 9.80% | 5.36% | 21.04% | 8.55% |
| pepper | 19.15% | 46.12% | 27.06% | 21.40% | 37.60% | 27.28% |

| Pacbio | | | | | | |
|---|---|---|---|---|---|---|
| clair3 | 88.62% | 94.31% | 91.38% | 83.63% | 87.09% | 85.32% |
| deepvariant | 82.59% | 91.57% | 86.85% | 77.36% | 83.52% | 80.32% |
| gatk4 | 30.98% | 84.81% | 45.38% | 35.46% | 78.16% | 48.79% |

***Table S4***
Benchmarking metrics (precision, recall and F1-score) were generated for SNPs and InDels using two validation sets: GIAB 4.2.1 and CMRG. The data corresponds to one lane/flowcell of illumina, 10X genomics, nanopore or PacBio sequencing of the Ashkenazim son (HG002).

| Regions | Variant Type | Zygosity | GIAB v4.2.1 | CMRG v1.0 |
|---|---|---|---|---|
| Easy-to-map | SNPs | Heterozyous | 1666055 | 5958 |
| | | Homozygous alt | 1052224 | 3837 |
| | | **Total** | 2718279 | 9795 |
| | InDels | Heterozyous | 92544 | 364 |
| | | Homozygous alt | 64205 | 267 |
| | | **Total** | 156749 | 631 |
| Difficult-to-map | SNPs | Heterozyous | 383620 | 2019 |
| | | Homozygous alt | 249707 | 1716 |
| | | **Total** | 633327 | 3735 |
| | InDels | Heterozyous | 201561 | 714 |
| | | Homozygous alt | 129740 | 554 |
| | | **Total** | 331301 | 1268 |

***Table S5***
Detailed variant counts dissecting GIAB's truth set: v 4.2.1 and CMRG under different genomic contexts, variant types and zygosity types.

| | **Deletions** | | | **Insertions** | | |
|---|---|---|---|---|---|---|
| Ref size ranges | All Tiers | Tier 1 | CMRG | All Tiers | Tier 1 | CMRG |
| [0,30) | 4413 | 0 | 0 | 4141 | 0 | 0 |
| [30, 50) | 3291 | 0 | 0 | 2604 | 0 | 0 |
| [50, 250) | 7444 | 2217 | 56 | 7164 | 2549 | 53 |
| [250, 500) | 2359 | 1181 | 19 | 2859 | 1535 | 27 |

| | | | | | | |
|---|---|---|---|---|---|---|
| [500, 6000) | 1682 | 611 | 14 | 3131 | 1074 | 25 |
| [6000, 260000000) | 225 | 108 | 0 | 190 | 103 | 0 |
| All ranges | 19414 | 4117 | 89 | 20089 | 5261 | 105 |

***Table S6***
Detailed stratification of SV deletion and insertion counts dissecting GIAB's SV truth set by Tier 1+2, Tier 1 amd CMRG in seven categories: 0-30 bp, 31-50 bp, 51-250 bp, 251-500 bp, 500-6000 bp and 6000-26000000 bp.

| Truth Set | Variant Type | F1 Type | Ave F1 10X | Ave F1 LRS | Ave F1 SRS | LRS_over_SRS | LRS_over_10X |
|---|---|---|---|---|---|---|---|
| cmrg | del | Genotyping F1 | 0.38 | 0.78 | 0.45 | 1.71 | 2.08 |
| cmrg | del | Discovery F1 | 0.59 | 0.94 | 0.63 | 1.51 | 1.60 |
| tier1 | del | Genotyping F1 | 0.50 | 0.95 | 0.72 | 1.32 | 1.91 |
| tier1 | del | Discovery F1 | 0.71 | 0.96 | 0.79 | 1.23 | 1.35 |
| tier1+2 | del | Genotyping F1 | 0.18 | 0.50 | 0.27 | 1.82 | 2.79 |
| tier1+2 | del | Discovery F1 | 0.40 | 0.73 | 0.43 | 1.68 | 1.84 |
| cmrg | ins | Genotyping F1 | *NA* | 0.68 | 0.22 | 3.17 | *NA* |
| cmrg | ins | Discovery F1 | *NA* | 0.87 | 0.33 | 2.63 | *NA* |
| tier1 | ins | Genotyping F1 | *NA* | 0.90 | 0.36 | 2.51 | *NA* |
| tier1 | ins | Discovery F1 | *NA* | 0.95 | 0.45 | 2.09 | *NA* |
| tier1+2 | ins | Genotyping F1 | *NA* | 0.51 | 0.17 | 3.03 | *NA* |
| tier1+2 | ins | Discovery F1 | *NA* | 0.74 | 0.29 | 2.60 | *NA* |

***Table S7***
Comparing the average discovery and genotyping F1 score across 1 10X (Longranger), 3 SRS (dragen, dysgu and metasv) and 4/5 LRS (CuteSV, dysgu, pbsv, sniffles2 and SVIM) SV callers and calculating the ratios between mean F1 score to compare technologies.

| | | | | | | **Ratio** |
|---|---|---|---|---|---|---|

| Truth Set | type | Ref size ranges | LRS | SRS | synthetic | LRS/SRS | LRS/synthetic | SRS/synthetic |
|---|---|---|---|---|---|---|---|---|
| cmrg | del | All ranges | 81.78 | 41.67 | 38.00 | 1.96 | 2.15 | 1.10 |
| cmrg | del | [250, 500) | 17.78 | 15.17 | 15.00 | 1.17 | 1.19 | 1.01 |
| cmrg | del | [50, 250) | 51.00 | 16.33 | 14.00 | 3.12 | 3.64 | 1.17 |
| cmrg | del | [500, 6000) | 13.00 | 10.17 | 9.00 | 1.28 | 1.44 | 1.13 |
| cmrg | del | [6000, 260000000) | 0.00 | 0.00 | 0.00 | *NA* | *NA* | *NA* |
| cmrg | ins | All ranges | 86.89 | 22.17 | *NA* | 3.92 | *NA* | *NA* |
| cmrg | ins | [250, 500) | 22.22 | 6.17 | *NA* | 3.60 | *NA* | *NA* |
| cmrg | ins | [50, 250) | 50.00 | 14.50 | *NA* | 3.45 | *NA* | *NA* |
| cmrg | ins | [500, 6000) | 14.67 | 1.50 | *NA* | 9.78 | *NA* | *NA* |
| cmrg | ins | [6000, 260000000) | 0.00 | 0.00 | *NA* | *NA* | *NA* | *NA* |
| tier1 | del | All ranges | 3902.67 | 2723.33 | 2485.00 | 1.43 | 1.57 | 1.10 |
| tier1 | del | [250, 500) | 1160.78 | 1034.50 | 1031.00 | 1.12 | 1.13 | 1.00 |
| tier1 | del | [50, 250) | 2105.44 | 1105.67 | 867.00 | 1.90 | 2.43 | 1.28 |
| tier1 | del | [500, 6000) | 548.89 | 498.67 | 503.00 | 1.10 | 1.09 | 0.99 |
| tier1 | del | [6000, 260000000) | 87.56 | 84.50 | 84.00 | 1.04 | 1.04 | 1.01 |
| tier1 | ins | All ranges | 4891.22 | 1583.33 | *NA* | 3.09 | *NA* | *NA* |
| tier1 | ins | [250, 500) | 1492.22 | 477.67 | *NA* | 3.12 | *NA* | *NA* |
| tier1 | ins | [50, 250) | 2389.00 | 962.67 | *NA* | 2.48 | *NA* | *NA* |
| tier1 | ins | [500, 6000) | 958.56 | 134.33 | *NA* | 7.14 | *NA* | *NA* |
| tier1 | ins | [6000, 260000000) | 51.44 | 8.67 | *NA* | 5.94 | *NA* | *NA* |
| tier1+2 | del | All ranges | 8898.22 | 4174.00 | 3740.00 | 2.13 | 2.38 | 1.12 |
| tier1+2 | del | [250, 500) | 1730.44 | 1245.17 | 1273.00 | 1.39 | 1.36 | 0.98 |
| tier1+2 | del | [30, 50) | 1358.78 | 282.00 | 122.00 | 4.82 | 11.14 | 2.31 |
| tier1+2 | del | [50, 250) | 4618.78 | 1807.50 | 1485.00 | 2.56 | 3.11 | 1.22 |
| tier1+2 | del | [500, 6000) | 1059.00 | 724.17 | 754.00 | 1.46 | 1.40 | 0.96 |
| tier1+2 | del | [6000, 260000000) | 131.22 | 115.17 | 106.00 | 1.14 | 1.24 | 1.09 |
| tier1+2 | ins | All ranges | 10676.56 | 2858.67 | *NA* | 3.73 | *NA* | *NA* |
| tier1+2 | ins | [250, 500) | 2242.00 | 704.67 | *NA* | 3.18 | *NA* | *NA* |
| tier1+2 | ins | [30, 50) | 1364.33 | 205.33 | *NA* | 6.64 | *NA* | *NA* |
| tier1+2 | ins | [50, 250) | 5122.56 | 1460.50 | *NA* | 3.51 | *NA* | *NA* |

| | | | | | | | | |
|---|---|---|---|---|---|---|---|---|
| tier1+2 | ins | [500, 6000) | 1873.22 | 446.17 | *NA* | 4.20 | *NA* | *NA* |
| tier1+2 | ins | [6000, 260000000) | 74.44 | 42.00 | *NA* | 1.77 | *NA* | *NA* |

***Table S8***

Comparing the average true positive (TP) counts across 1 10X (Longranger), 3 SRS (dragen, dysgu and metasv) and 4/5 LRS (CuteSV, dysgu, pbsv, sniffles2 and SVIM) SV callers for each seven categories: 0-30 bp, 31-50 bp, 51-250 bp, 251-500 bp, 500-6000 bp and 6000-26000000 bp then calculating the ratios between mean TP score to compare technologies.
